# Global proteomics indicates subcellular-specific anti-ferroptotic responses to ionizing radiation

**DOI:** 10.1101/2024.09.12.611851

**Authors:** Josie A. Christopher, Lisa M. Breckels, Oliver M. Crook, Mercedes Vazquez--Chantada, Derek Barratt, Kathryn S. Lilley

## Abstract

Cells have many protective mechanisms against background levels of ionizing radiation (IR) orchestrated by molecular changes in expression, post-translation modifications and subcellular localization. Radiotherapeutic treatment in oncology attempts to overwhelm such mechanisms, but radio-resistance is an ongoing challenge. Here, global subcellular proteomics combined with Bayesian modelling identified 544 differentially localized proteins in A549 cells upon 6 Gy x-ray exposure, revealing subcellular-specific changes of proteins involved in ferroptosis, an iron-dependent cell death, suggestive of potential radio-resistance mechanisms. These observations were independent of expression changes, emphasizing the utility of global subcellular proteomics and the promising prospect of ferroptosis-inducing therapies for combatting radioresistance.

## Introduction

Radiotherapy, or ionizing radiation (IR), has been a long-standing curative and palliative treatment option for cancers, as well as other diseases. The high energy of IR causes a cascade of lethal biochemical reactions within cells, initiated by the ionization of water and production of free radicals, causing double-stranded breaks (DSBs) in DNA, lipid peroxidation and resulting tissue damage (De Ruysscher et al., 2019). Whilst IR toxicity to healthy tissue is a concern, higher radiosensitivity of actively dividing and undifferentiated cells versus mature and non-dividing cells allows a sufficient therapeutic window for IR to be an effective cancer treatment (De Ruysscher et al., 2019; Meyniel, 1958). More precise delivery methods for IR, such as stereotactic ablative radiotherapy (SABR) (Shenker et al., 2022), have increased the therapeutic index between tumor and normal tissue response, allowing for increased doses with reduced side effects. These modern approaches have prevented IR from becoming obsolete and being superseded by molecular target-based pharmaceuticals, with approximately 50% of cancers still using radiotherapy as a component of their treatment regime (Howlader et al., 2013; Xing and Stea, 2024). However, naturally occurring background radiation in the environment causes a continuous, chronic risk to cellular DNA. Thus, cells have evolved accordingly to have multiple DNA damage repair and antioxidant pathways to subdue the cascading effect of IR-induced free radicals (Espinosa-Diez et al., 2015). Therefore, as with many other oncotherapeutics, resistance to radiotherapy can be a significant issue (Xing and Stea, 2024).

In an attempt to improve efficacy of treatment, radiotherapy is commonly combined with therapeutics that target auxiliary pathways. For example, as the predominant mechanism of action of IR is DNA damage, there is interest in combining radiotherapy with inhibitors targeting DNA damage response pathways, such as DNA repair and cell cycle checkpoints (Morgan and Lawrence, 2015). Targeting of the DNA repair pathways in cancer alone is also of high interest, as cancers are characteristically genetically unstable and generally have impaired DNA repair (e.g. BRCA and ATM mutants), and thus rely more heavily on alternative repair pathways to survive during chemotherapy (O’Connor, 2015). Therefore, studying response mechanisms to IR is well placed to elucidate the mechanisms by which tumor cells avoid cell death from DNA-damaging agents.

Proteomics approaches have rapidly accelerated our understanding of cellular biology and aided drug discovery with untargeted, high-throughput screening (Nguyen et al., 2022). However, generally these approaches focus on changes in protein expression and lack spatial information - i.e. where a molecule resides within a cell, its surrounding microenvironment and proximity to molecular interactors. This potentially overlooks key mechanisms in cellular functions, responses, and pathological mechanisms. For example, many diseases have been associated with defective protein trafficking, such as cystic fibrosis, amyotrophic lateral sclerosis and pulmonary atrial hypertension, which may have unremarkable expression profiles of the affected proteins (Cheng et al., 1990; Christopher et al., 2021; Guo and Shorter, 2017; Ren et al., 2013, p. 20113; Sehgal and Lee, 2011). Whilst microscopy provides such information, this technology is generally reserved for targeted experiments and suffers from limitations with gene fusion artefacts or non-specific antibodies (Baker, 2015; Stadler et al., 2013). Proximity labelling approaches, where a protein of interest is genetically fused with a promiscuous enzyme capable of generating activated biotin that labels its immediate subcellular environment, has also been used to map subcellular distributions of proteins (Go et al., 2021). This approach is difficult to scale to a cell-wide method, requires genetic modification and also suffers from differential labelling in different subcellular niches leading to a biased map of subcellular residency of proteins (Christopher et al., 2021). This study used an untargeted, subcellular proteomics method, Localization of Proteins using Isobaric Tagging using Differential Centrifugation (LOPIT-DC) (Braccia et al., 2022; Geladaki et al., 2019), to capture subcellular information of thousands of proteins in multiple subcellular compartments before and after 6 Gy IR in the A549 lung carcinoma cell line to detect trafficking-specific responses to IR. Combining this untargeted, unbiased cell-wide approach with targeted validation, several subcellular-specific changes of proteins associated with the iron-dependent regulated cell death pathway, ferroptosis, were detected. This indicated a radio-resistant response to IR within these cells, alongside expression changes of related pathways.

## Experimental Procedures

### Cell culture & x-ray treatment

Lung epithelial carcinoma cell line, A549 (CCL-185^TM^, ATCC®) were certified as mycoplasma-free. Cells were not kept in culture for longer than a month and were grown in Ham’s F12 (Sigma Aldrich), with 10% fetal bovine serum (FBS, ThermoFisher, Lot: 42F3393K) and 1% L-glutamine. Cells were passaged at around 70% confluence by washing with PBS before dissociating from the flask with TrypLE^TM^ Express (Gibco^TM^, 12604013) and kept at 37°C and 5% CO_2_. Total absorbed dose of 6 Gy using either a Faxitron® CellRad® or Pantak HF320kV x-ray system. The Pantak HF320kV x-ray system operated at 220 kV, 14 mA with a 0.5 Cu filter with a dose rate of 428 cGy/min. The Faxitron® CellRad® using a 50 Cu with a kV/mA dependent on the shelf used on the instrument. Control cells were sham irradiated (kept in the same conditions as the IR-treated cells).

### Localization of Proteins using Isobaric Tagging using Differential Centrifugation (LOPIT-DC)

Performed as previously published (Geladaki et al., 2019). Briefly, cells were harvested, washed and resuspended in an isotonic lysis buffer 12 hrs after (sham) irradiation. Cell plasma membrane (PM) lysis was performed using a ball-bearing homogenizer (Isobiotec) using a 12 µM clearance. Differential centrifugation was performed using a Beckman Optima MAX-XP ultracentrifuge (with a TLA-55, fixed angle rotor) with spin speeds and times in Table S1. Samples were always kept on ice or at 4°C throughout the procedure. The resulting supernatant was precipitated using chilled acetone. Organellar-enriched pellets were solubilized using a urea-based buffer and sonication. Organellar enriched lysates then underwent reduction, alkylation, digestion, TMT labelling, clean-up and LC-MS/MS as documented below. (Fig. 1, Detailed in Supplementary Methods.)

**Figure 1.**
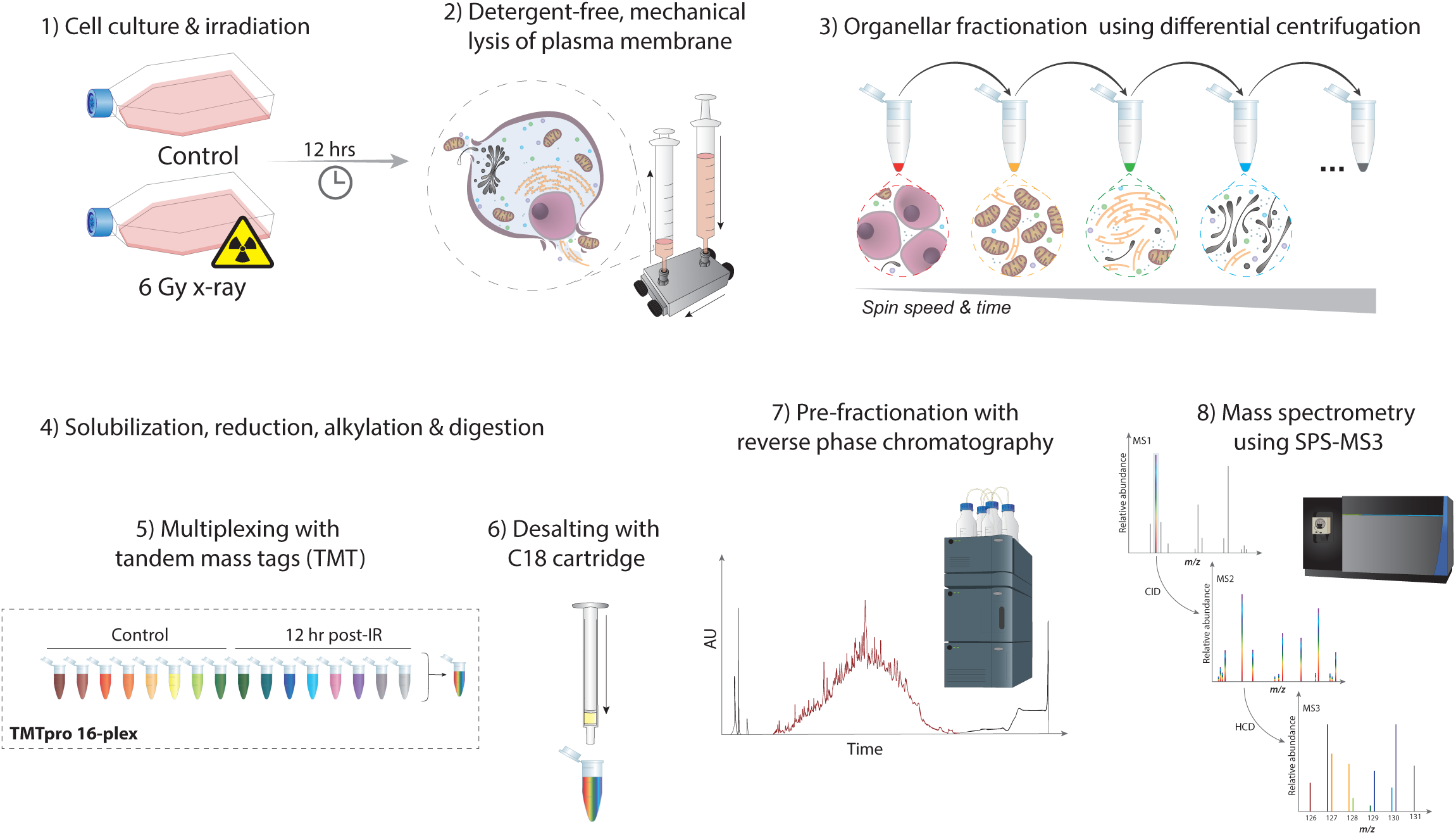
LOPIT-DC sample preparation and LC-MS/MS workflow. Briefly, A549 cells were either dosed with 6 Gy X-rays or sham-irradiated (control) (1), then 12 hrs later plasma membrane (PM) lysed using a ball-bearing homogeniser (2). Organellar enrichmed pellets were achieved using differential centrifugation using a gradient of different spin speeds and time (3) before pellet solubilisation using urea-based buffer and sonication followed by dithiothreitol (DTT) reduction, iodoacetamide (IAA) alkylation and trypsin digestion (4). Solubilised pellets were labelled with tandem mass tags (TMT) (5), pooled, and HEPES removed using C18 desalting cartridges (6). Offline fractionation of the pooled TMT-labelled organellar-enriched sample performed using reverse-phase chromatography (7) before analysing using an Orbitrap mass spectrometer in SPS-MS3 mode (8).

### Total proteomics harvesting, protein digestion, TMT labelling & clean-up

Cells were harvested and washed 12 hrs after (sham) irradiation, as LOPIT-DC experimental procedure, and lysed in a urea-based buffer using sonication. LOPIT-DC and total proteomics samples were processed in the same manner from this point on. Protein concentrations of LOPIT-DC pellets and total proteomics samples were measured using BCA protein assay and were normalized to 50 μg/100 μL lysis buffer. Samples underwent dithiothreitol (DTT) reduction, iodoacetamide (IAA) alkylation and acetone precipitated, followed by trypsin digestion at 37°C. Tandem mass tag (TMT) labelling was performed as per the manufacturer’s instructions. TMT 10-plex (ThermoFisher, 90406) were used for total proteomics sample labelling and TMTpro 16-plex (ThermoFisher, A44520) were used for LOPIT-DC pellet labelling. Samples were lyophilized before performing solid phase extraction using SepPak® C18 columns (Waters, WAT054955). Peptides were washed with 0.1% TFA, eluted with 70% ACN/0.05% acetic acid and lyophilized. (Detailed in Supplementary Methods.)

### LC-MS/MS

Total proteomics and LOPIT-DC samples were pre-fractionated offline using reverse phase UPLC with an Acquity UPLC System with diode array detector (Waters) and an Acquity UPLC BEH 283 C18 column (2.1-mm ID × 150-mm; 1.7-µm particle size) (Water, 186002353). Peptide fractions were collected every minute of the gradient over a 50 min linear gradient resulting in 15 concatenated fractions. Samples were lyophilized and resuspended in 0.1% formic acid. MS analysis was performed as previously published (Mulvey et al., 2017), and detailed in Supplementary Methods.

### Proteomics data processing & analysis

Raw MS files were processed using Proteome Discoverer 2.4 (ThermoFisher) and a Mascot server (Matrix Science) v2.6 (Brosch et al., 2009). SwissProt Homo sapiens database (no isoforms, 20,245 sequences, downloaded 09/04/2018) and the common repository of adventitious proteins (cRAP, v1.0, 123 sequences) were used. Fixed modifications were set to carbamidomethyl(C) and TMT 10-plex or TMTpro(K, peptide N termini), and variable modifications were set to carbamyl(N-term), carbamyl(K), carbamyl(R) and deamidation(N,Q). (Further detailed in Supplementary Methods.)

Proteomics data was analyzed in R (v4.1.3) using the limma (v3.50.3) Bioconductor package (Ritchie et al., 2015). An empirical Bayes moderated t-test was performed for differential expression analysis of the total proteomics data (Smyth, 2004). P-values were adjusted for multiple testing using the Benjamini Hochberg method (Benjamini and Hochberg, 1995). Gene ontology (GO) enrichment analysis was performed by using the R package, clusterProfiler (v4.2.2) (Gene Ontology Consortium, 2021; Wu et al., 2021). Analysis of LOPIT-DC used the spatial proteomics package, pRoloc (v1.34.0) (Gatto et al., 2014). A semi-supervised machine learning approach, Bayesian ANalysis of Differential Localization Experiments (BANDLE), was used to classify proteins to subcellular localization, identify differential localization, and estimate uncertainty of those predictions (Crook et al., 2021). Subcellular protein markers for this algorithm were established using existing curated markers (Geladaki et al., 2019; Mulvey et al., 2017; Orre et al., 2019), the bioinformatics tool COMPARTMENTS (Binder et al., 2014), and the unsupervised, clustering algorithm, HDBscan (Melvin et al., 2016), plus other filtering criteria documented in the Supplementary Methods. BANDLE analysis was performed as documented in the Bioconductor vignettes (https://bioconductor.org/packages/release/bioc/html/bandle.html). (Detailed in Supplementary Methods.)

### Flow cytometry

Manufacturer protocols for propidium iodide (Abcam, ab139418) and CellEvent™ Caspase-3/7 (Invitrogen^TM^, C10427) staining were followed for the cell cycle and cell death assays, respectively. Cells were treated with 1 µg/ml nocodazole for 18 hrs (as previously published (Queiroz et al., 2019)) or 100 μg/mL sodium arsenate for 2 hrs as positive controls for cell arrest and cell death, respectively. Sodium arsenate was also added to compensation samples. Cells were analyzed on an Attune™ NxT Acoustic Focusing Cytometer.

Flow cytometry analysis was performed using the FCS ExpressTM software (De Novo Software^TM^, version 7.12). Gates were set to exclude debris and doublets from the analysis. Cell cycle analysis was performed using the MultiCycle algorithm feature in the FCS Express software, using the Dean & Jett mathematical model (Dean and Jett, 1974)). Compensation matrixes were set using the compensation control samples. Statistical analysis and visualization for both the cell cycle and cell death assay results were performed in R using a pair-wise t-test and Bonferroni adjustment. (Detailed in Supplementary Methods.)

### Immunofluorescence microscopy

Cells were seeded into a glass-bottomed 8-high-well µ-Slide (Ibidi, 80806), exposed to 6 Gy x-ray and incubated for 12 hrs. Cells were stained with MemBrite® Fix 640/60 cell surface stain (Biotium, 30097) prior to fixation and according to manufacturer’s guidelines. Cells were then fixed with 4% paraformaldehyde (PFA) and permeabilized using 0.1% Triton X-100. Primary antibodies were added at the manufacturer or optimized concentration and incubated overnight at 4°C. Cells were washed and then incubated with corresponding IgG (H+L) conjugated with Alexa Fluor 568 (Invitrogen, A11004) or Alexa Fluor 488 (Invitrogen, A-11008) with 1:2,000 dilution before adding ProLong™ Gold Antifade Mountant with DAPI (Invitrogen, P36931). Single-stained and secondary only controls were prepared to assess non-specific binding or autofluorescence. See Table S2 for antibody details.

A Zeiss LSM 880 confocal microscope with a 40x/1.30 Plan Apo Oil objective lens and a Piezo stage was used for image acquisition. Fluorophores were excited using 405 nm (blue diode), 488 nm (Argon), 561 nm (He 543) and 633 nm (He 633) lasers with 0.05%, 1.5% and 3.0% power, respectively. Signal was detected using two photomultiplier (PMT) and a gallium arsenide phosphide (GaAsP) array detector in combination with filters set to 400-440 nm, 507-552 nm, 585-620 nm, 650-690 nm, to detect DAPI, AlexaFluor488, AlexaFluor568, and MemBrite® Fix 640/60, respectively. (Detailed in Supplementary Methods.) Colocalization analysis, segmentation and quantification of nuclear- and cytosolic-bound γH2AX foci was performed using the open-source CellProfiler software (v4.2.1 Broad Institute) (Stirling et al., 2021). (Detailed in Supplementary Methods.)

## Results

### DSBs and total proteome level changes were detected without marked changes in cell cycle or cell death 12hrs post-IR

At 12 hrs post-IR double-stranded breaks (DSBs) were confirmed by immuno-probing for phosphorylated serine 139 on histone H2AX (γH2AX) and quantitative imaging for γH2AX foci in the nuclei (Fig. 2A-B, S1). Cytosolic γH2AX foci measurements were also taken as a control. These data demonstrated that these cells responded typically to x-ray exposure (Bouquet et al., 2006). Additionally, total proteomics further confirmed biological response to x-ray exposure with dominant variance in principal component analysis (PCA) driven by control versus IR-exposed experimental groups (Fig. 2C). Flow cytometric measurements did not detect significant changes in cell cycle and cell death populations 12 hrs post-IR (Fig. 2D).

**Figure 2.**
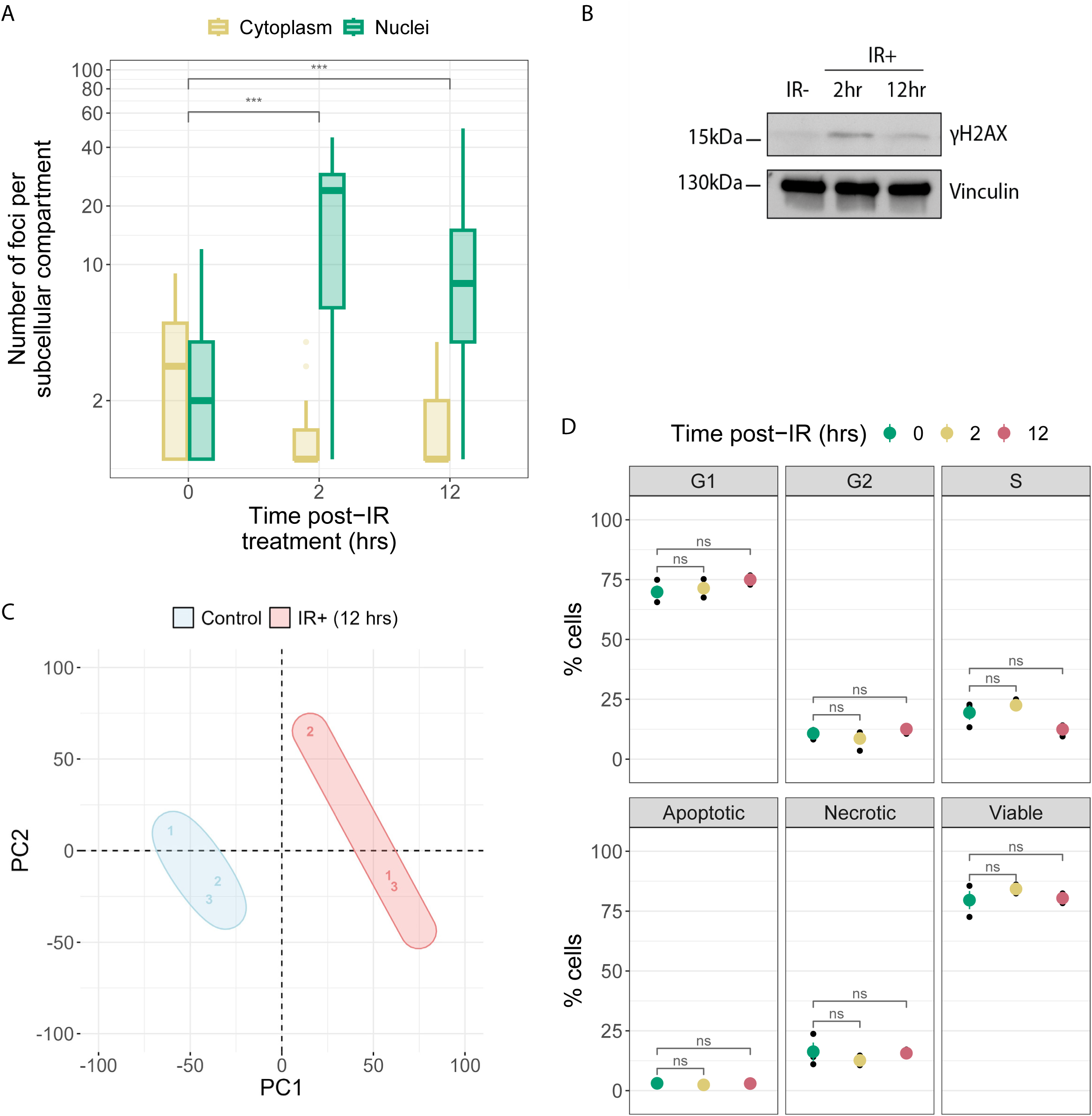
DNA damage, cell cycle status, apoptotic status and proteome-level changes 2 and 12 hrs after IR exposure. (A) Number of yH2AX foci measured in the nuclei and cytoplasm 2 and 12 hrs post-IR exposure using quantitative confocal microscopy, (B) with corresponding western blot probing yH2AX at these time points. (C) Principle component analysis (PCA) of total proteomics data for control or at 12 hrs post-IR. (D) Flow cytometry propidium iodide and CellEvent^TM^ Caspase3/7 assays measuring cell cycle (top) and apoptotic status (bottom), respectively, at 0, 2, and 12 hrs post-IR.

### Upregulated proteins identify characteristic cellular responses to IR

A total of 6,163 unique proteins were quantified in the total proteomics dataset, with 101 and 142 proteins up- and down-regulated post-IR (FDR < 0.01), respectively (Fig. 3A, Supplementary Data 1). Surprisingly, many REACTOME (Milacic et al., 2024) and cellular compartment terms that are enriched in the downregulated proteins are related to DNA repair, which may have expected to be upregulated after IR (Fig. 3B, S2). However, many of the proteins that are involved in DNA damage repair are also involved in cell cycle progression, which is reduced or halted after IR exposure. While this observation contrasts with what was seen in the above flow cytometry analysis, this suggests proteomic-level detection of initiation of cell cycle arrest mechanisms may precede the morphological and physiological changes measured in flow cytometry (i.e. differences in DNA content in cell cycle stages) (Fig. 2D). Phosphoproteomic analysis on these samples showed increased phosphorylation of key DNA damage repair proteins at this time point post-IR (Fig. S2, Supplementary Data 2, Supplementary Methods), demonstrating the nuances of molecular signaling of DNA repair in early cellular response to IR-induced DNA damage.

**Figure 3.**
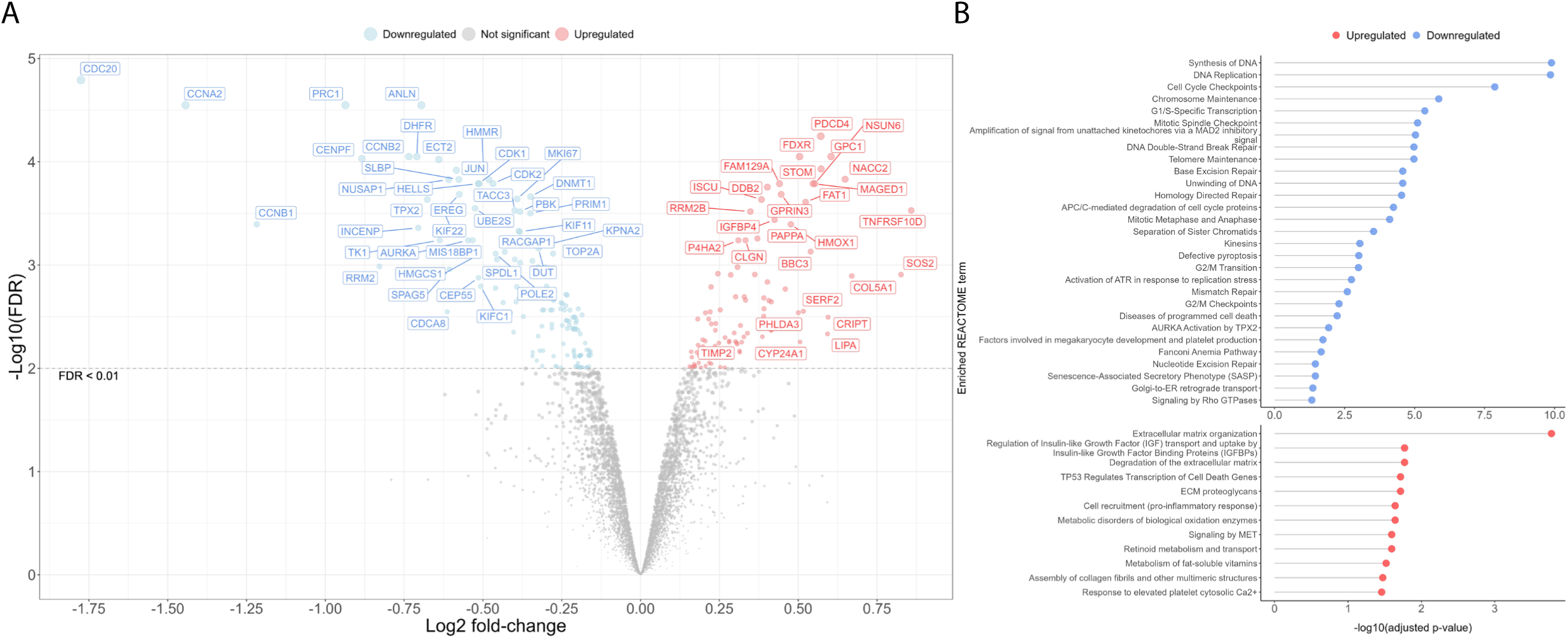
Total proteomics data for control versus IR-treated A549 cells 12 hrs post-IR. (A) Volcano plot of differential expression analysis for the total proteomics data in control verus 12 hrs post-IR in A549 cells. Coloured points denote all proteins with FDR < 0.01 and all labelled proteins with FDR < 0.001. (B) Enriched REACTOME terms for down- and up-regulated proteins (FDR < 0.01) 12 hrs post-IR.

Upregulated proteins were enriched for REACTOME terms, such as extracellular matrix (ECM) and retinoid metabolism. Previous studies have demonstrated that retinoids have a radioprotective effect, preventing radiation-induced apoptosis in cells (Cho et al., 2010; Vorotnikova et al., 2004) and have been found to upregulate tissue inhibitors of matrix metalloproteinases (TIMPs) (Vorotnikova et al., 2004). Upregulation of TIMP2, alongside other proteins involved in ECM organisation such as collagens, laminins and integrins, was detected in the total proteomics analysis (Fig. 3A-B). Prominent upregulation of these proteins is unsurprising, as IR-related dermatitis can be induced at levels of 6 Gy or more (Kole et al., 2017). These changes in the ECM are also responsible for much of the downstream signaling and biological responses to this IR-induced damage, such as MET signaling, which was also an enriched pathway (Fig. 3B). MET is essential for wound healing and activation of MAPK and PI3K-Akt signal transduction (Trusolino et al., 2010), which participate in DNA damage, oxidative stress responses, and cell fate (Karimian et al., 2019; Toulany and Rodemann, 2015). Additionally, upregulated proteins involved in insulin growth factor (IGF) signaling were also detected and blockade of IGF signaling has shown to radiosensitize cells and be linked to radiation-mediated regulation of Ku86 via p38 (Cosaceanu et al., 2007; Y Aghdam et al., 2021). Interestingly, an enriched cellular component was membrane (or lipid rafts), sphingolipid-rich membrane domains that compartmentalize signaling processes within the cell (Fig. S3). Pro-survival signaling pathways such as PI3K-Akt and IGF signaling, which cancer cells are known to hijack, is known to be modulated via lipid rafts (Mollinedo and Gajate, 2020).

### LOPIT-DC captures the dynamics of the subcellular proteome post-IR

LOPIT-DC datasets were collected in biological triplicate and the resulting data processed as in Geladaki et al., 2019 ((Geladaki et al., 2019) Supplementary Data 3). Approximately 85% of proteins quantified in each LOPIT-DC sample were detected across all replicates, with 5,785 and 5,773 proteins found in control and IR-treated samples, respectively, and 5,767 proteins shared across all LOPIT-DC samples (Fig. 4A, S4). The consistency of the protein identifications, correlation profiles and principal component analysis (PCA) demonstrated the high reproducibility of these experiments (Fig. S5, S6A-B). A semi-supervised machine learning approach, BANDLE, with 1,509 manually- and computationally-curated subcellular protein markers from previous studies, databases and literature (Fig. 4D) classified 1,913 and 2,179 proteins to 12 subcellular classes within the control and IR-treated groups, respectively (Fig. 4E). BANDLE was also used to calculate which proteins had differentially localized post-IR exposure. The term “differentially localized” is to describe proteins that appear to have changed localization after stimuli based on LOPIT-DC analysis, though this does not rule out that a protein has changed in abundance in one compartment over another. Therefore, the term “trafficking” or “translocation” is avoided.

**Figure 4.**
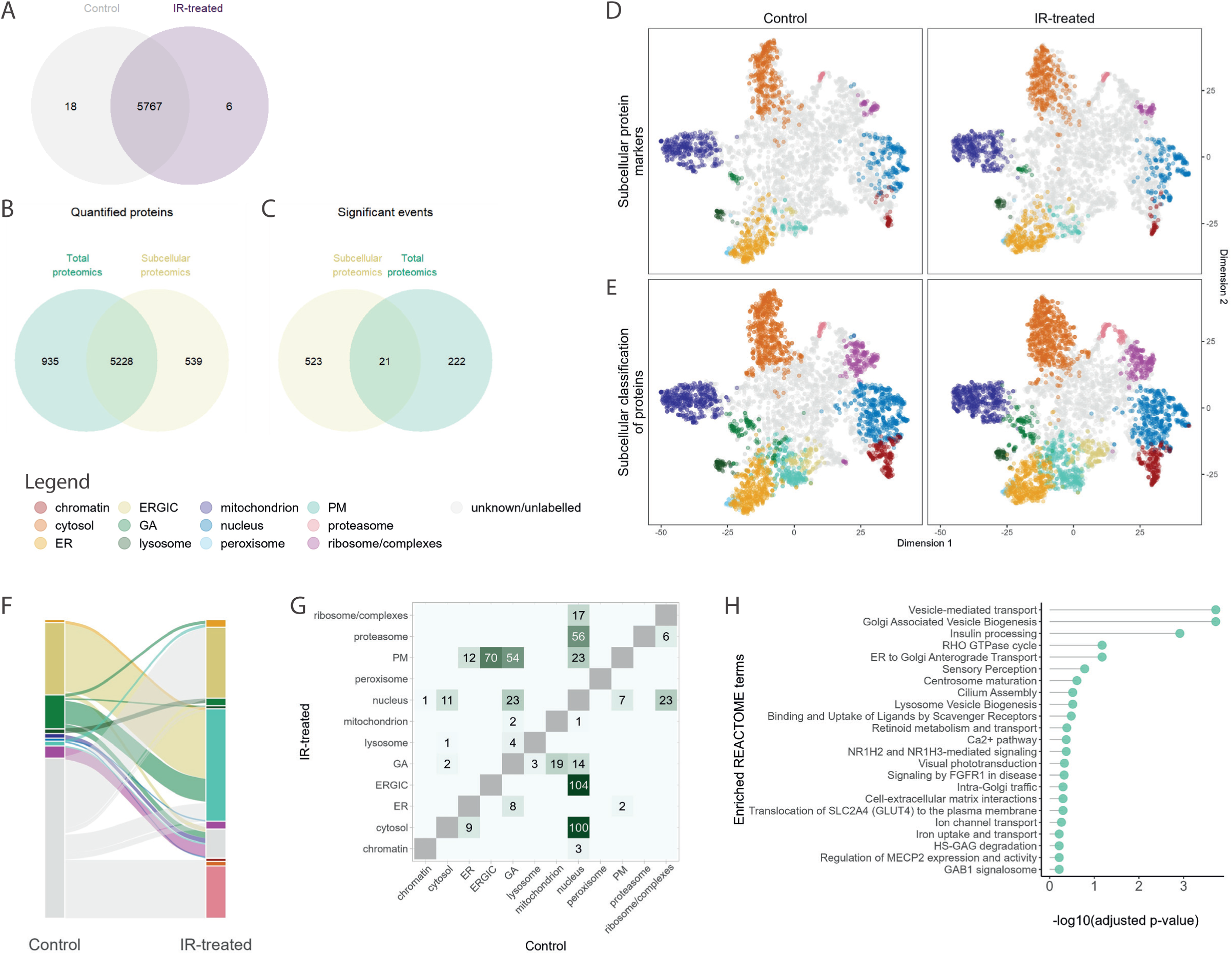
Subcellular proteomics using LOPIT-DC to identify changes in cellular location of proteins 12 hrs after IR exposure. (A) Venn diagrams of shared protein identifications between the control and IR-treated groups in LOPIT-DC experiment, (B) the quantified protein identifications in the total proteomics data versus subcellular proteomics data, and (C) proteins that were considered significant in the LOPIT-DC data (eFDR < 0.01) and the total proteomics data (FDR < 0.01). (D) Combined t-distributed Stochastic Neighbor Embedding (t-SNE) projections of the LOPIT-DC control and IR-treated replicates, where each point represents an individual protein identification. Each colour denotes the protein markers of 12 distinct subcellular compartments and (E) the subcellular classification of proteins using these protein markers with BANDLE. (F) Sanky plot representing the 544 proteins identified as differentially localised after IR exposure by BANDLE, and (G) corresponding heatmap with numbers representing number of proteins localising between these subcellular compartment between control and IR-treated groups. Heatmap shows the subcellular classification irrespective of whether the classification was considered an outlier/”unknown” classification. (H) Enriched REACTOME terms for proteins identified as differentially localising. (ER, endoplasmic reticulum; ERGIC, endoplasmic-reticulum–Golgi intermediate compartment; GA, Golgi apparatus; PM, plasma membrane).

BANDLE also identified 544 differentially localized proteins (estimated false discovery rate (eFDR) < 0.05) (Fig. 4F). The majority of the proteins that were identified as differentially localizing are transported between organelles linked to the secretory pathway and cytoskeletal remodeling, with enriched REACTOME terms such as “Golgi associated vesicle biogenesis” and “RHO GTPase”, respectively (Fig. 4G-H). It is worth noting that the Golgi apparatus (GA) is known to fragment or disperse in response to IR, and is even thought this fragmentation participates in DNA repair signaling via DNA-PKcs, increasing cell survival (Farber-Katz et al., 2014; Oike et al., 2021). These data also suggest increased vesicle transport to the PM. For example, exocyst complex proteins, which participate in tethering secretory vesicles to the PM, differentially localized to the PM post-IR (Fig. S7) (Mei and Guo, 2018). Terms that are non-specific to the secretory pathway and protein trafficking were also enriched, such as “binding and uptake of ligands by scavenger receptors”, “metabolism of fat-soluble vitamins”, “iron uptake and transport” and “transferrin endocytosis and recycling” (Fig. 4H). Interestingly, a lot of these terms and the proteins themselves are linked to lipid and iron metabolism, indicating subcellular-driven responses to IR-induced lipid peroxidation (Ye et al., 2020; Yu et al., 2015).

### Little overlap between differentially localized and differentially expressed proteins

Of the 5,228 proteins that were quantified in both the LOPIT-DC and the total proteomics experiments, only 21 proteins were found to be differentially expressing (FDR < 0.01) and localizing (eFDR < 0.05) after IR (Fig. 4B-C). This suggests that 96% of the proteins that were considered to have differentially localized (eFDR < 0.05) were likely protein trafficking events, rather than driven by a change in abundance and accumulation of a protein in a particular subcellular compartment. However, at this time point, this could also represent newly synthesized proteins that have trafficked to a different localization to where their older, degraded counterparts were predominantly localized. While differentially regulated proteins largely differed between the total proteomics and LOPIT-DC experiments, similar or connected REACTOME terms were enriched between these datasets (Fig. 3B and 4H). Proteins from the same or similar REACTOME pathways were detected as changing in expression or localization but infrequently both, demonstrating the complementary nature of using these two different untargeted proteomics methods (Fig. S9). Proteins upregulated in the total proteomics data had overlapping terms to the differentially localized proteins, which included lipid and retinoid metabolism, as well as pathways linked to insulin-like growth factor (IGF) signaling and ECM organization, such as GAB1 signaling and HS-GAG signaling, respectively (Fig. 3B and 4H) (Griffin and Gloster, 2017; Winnay et al., 2000). These biological processes and pathways are all highly interconnecting and biologically relevant to IR damage and response.

### LOPIT-DC detected subcellular-specific changes to key ferroptosis proteins

In addition to the above observations relating to lipid metabolism, retinoid metabolism and scavenger receptors, subcellular changes to several key proteins that are part of the iron-dependent cell death pathway, ferroptosis were observed (Bi et al., 2023; Chen et al., 2021). These proteins included transferrin receptor C (TFRC), the heavy and light chains of ferritin (FTH1 and FTL), and apoptosis-inducing factor homologous mitochondrion-associated inducer of death 2 (AIFM2) (Fig. 5A-B). However, in recent years AIFM2 has been renamed ferroptosis suppressor protein (FSP1) due to its significant role in glutathione-independent inhibition of ferroptosis (Bersuker et al., 2019; Doll et al., 2019). AIFM2, TFRC and FTL did not change in total abundance according to the total proteomics data, suggesting probable protein trafficking events (Fig. 5C). FTH1 was not detected in the total proteomics analysis. The subcellular localization of these proteins was verified using confocal microscopy (Fig. 6).

**Figure 5.**
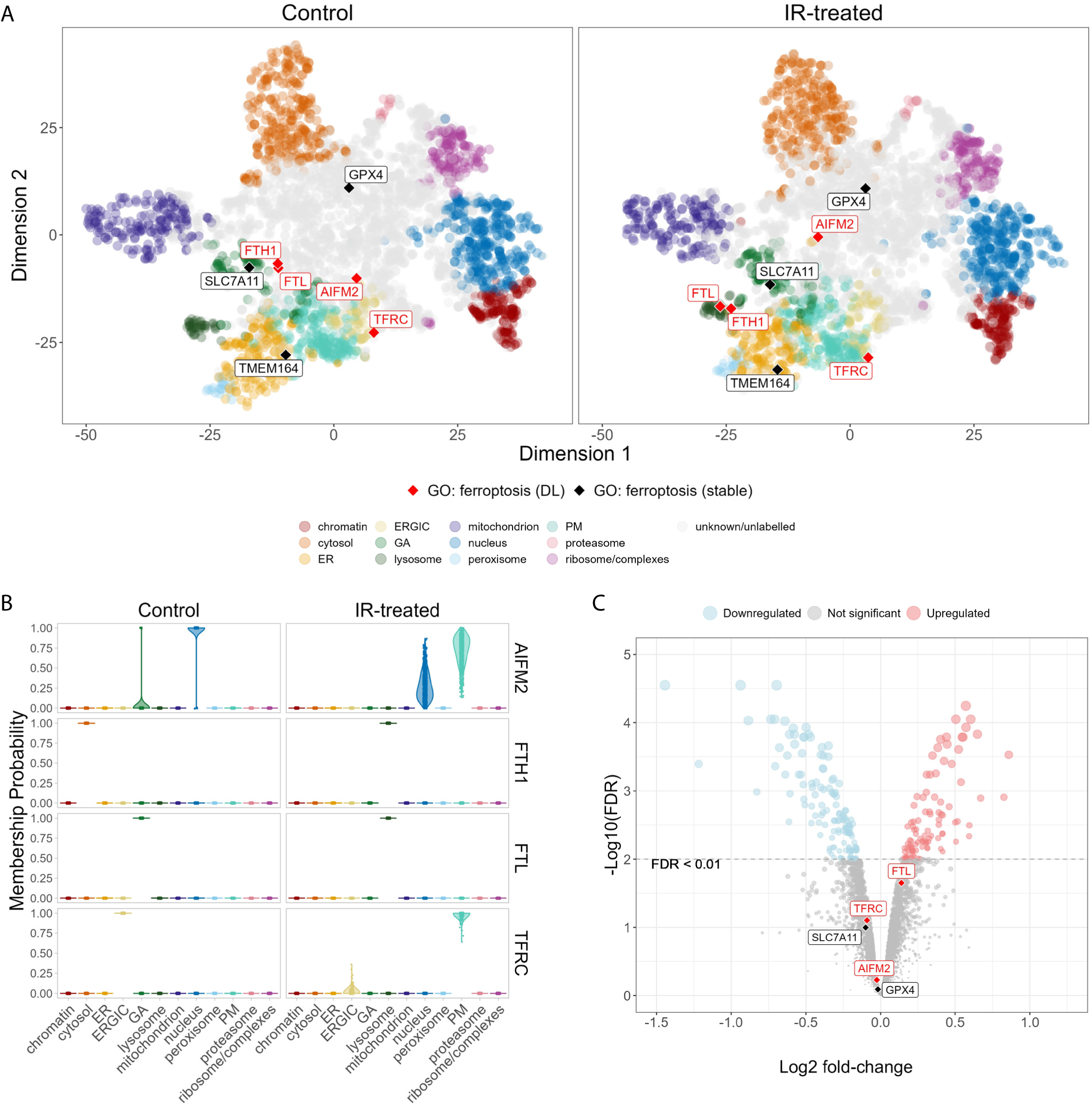
Ferroptosis proteins in the subcellular and total proteomics data. (A) t-SNE projections of control vs IR-treated labelled with proteins with associate gene ontology (GO) term “ferroptosis”. Red and black labels are proteins that were either identified as differentially localised or stable after IR by BANDLE (eFDR < 0.01). (B) Posterior predictive distributions (membership probability) across the 12 subceullar compartments for proteins identified as differentially localised and part of ferroptosis pathway in control vs IR-treated LOPIT-DC samples. (C) Volcano plot annotated with ferrotosis proteins that were measured in the total proteomics data. (ER, endoplasmic reticulum; ERGIC, endoplasmic-reticulum–Golgi intermediate compartment; GA, Golgi apparatus; PM, plasma membrane).

**Figure 6.**
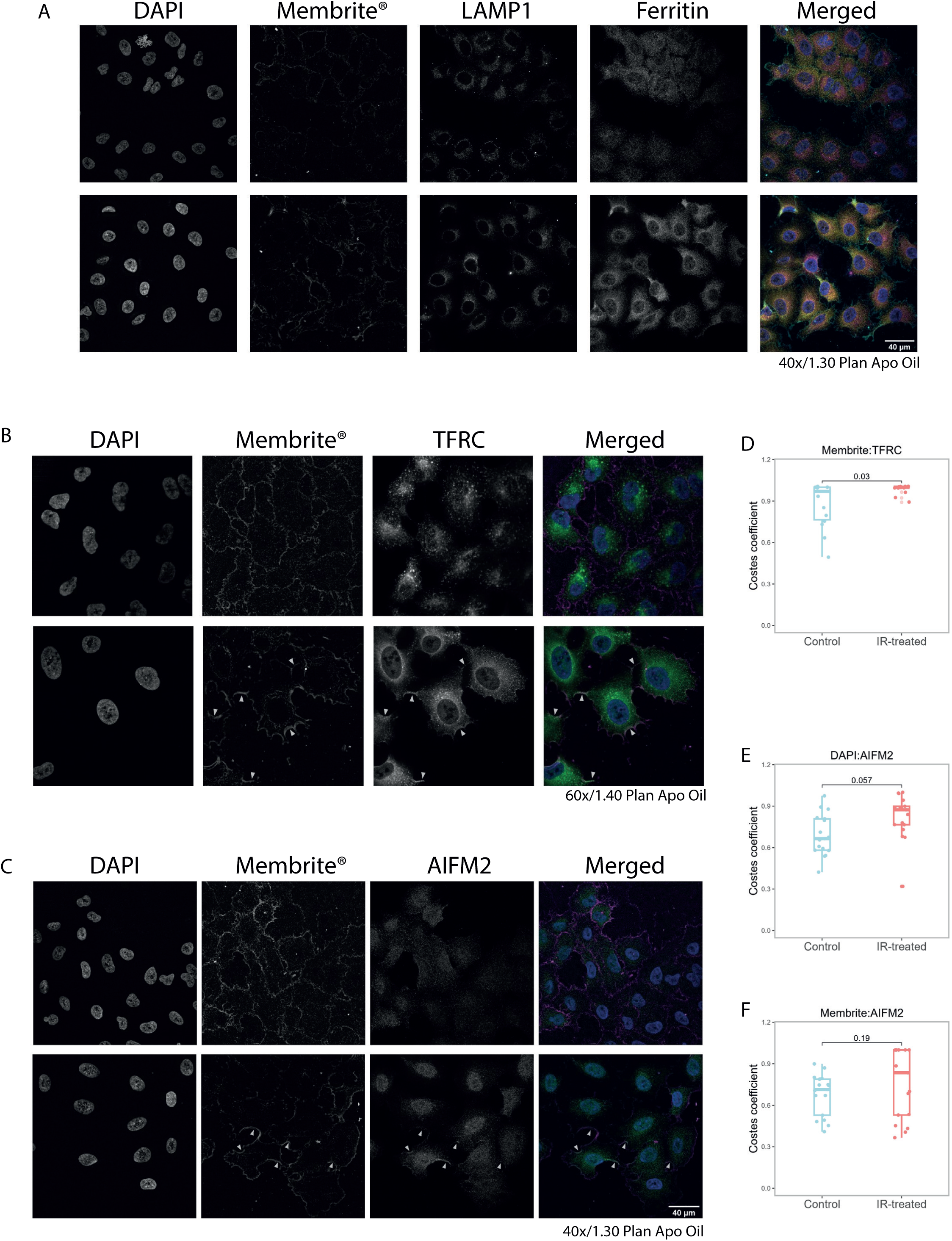
Validation of subcellular localisation of ferroptotic protein using confocal microscopy in control versus 12 hrs post-IR. Confocal images of A549 cells control vs 12 hrs post-IR with immunostaining of (A) ferritin (yellow), with lysosomal LAMP1 (magenta), and PM Membrite® stain (cyan); (B) TFRC (green), with PM Membrite® stain (magenta); (C) AIFM2 (green), with PM Membrite® stain (magenta). All images use nucleic DAPI staining (blue). Corresponding quantitative microscopy analysis to assess co-localisation of (D) Membrite® and TFRC, (E) DAPI and AIFM2, and (F) Membrite® and AIFM2.

Within the LOPIT-DC data, AIFM2 was classified as “unknown” in both the control and IR-treated groups, suggesting a multi-localized distribution within the cell (Fig. 5A). However, application of the BANDLE algorithm showed a shift of steady-state localization from a nuclear/Golgi profile to a more PM/nuclear profile (Fig. 5B). AIFM2’s primary, unperturbed steady-state localization has previously been observed in the Golgi (Varecha et al., 2007) and demonstrated to exert its ferroptosis suppressor function via PM and lipid droplets-targeted translocation (Bersuker et al., 2019). Interestingly, two proteins that are part of the glutathione-dependent ferroptosis inhibition pathway, phospholipid hydroperoxide glutathione peroxidase (GPX4) and cystine/glutamate transporter (SLC7A11) (Seibt et al., 2019), appeared to remain static and lack changes in expression post-IR (Fig. 5A,C, S8A). This combination of observations suggest A549 cells have a preference for glutathione-independent suppression of ferroptosis to avoid cell death upon IR exposure. Subcellular classification of AIFM2 was validated using confocal imaging and quantitative co-localization analysis comparing AIFM2 staining to nucleic DAPI and PM Membrite® staining. Control-AIFM2 was observed to be dispersed throughout the cells and then localized more discreetly to the nuclei and PM post-IR (Fig. 6C). Despite p-values above 0.05, small increases in quantitative co-localization of AIFM2 with Membrite® and DAPI post-IR were measured, with median Costes coefficient increased by 0.208 (p-value 0.057) and 0.121 (p-value 0.19), respectively (Fig. 5E-F). Whilst currently in contention due to the novel role of AIFM2 in ferroptosis inhibition, nuclear AIFM2 has been previously linked to caspase-independent apoptosis (Ohiro et al., 2002; Zheng and Conrad, 2020). Specifically, adduction of the lipid peroxidation product, 4-hydroxy-2-nonenal (4-HNE), induced AIFM2 nuclear translocation from the mitochondria to the nuclei as part of pro-apoptotic signaling (Miriyala et al., 2016). This evidence suggests A549 cells’ preference for caspase-independent apoptosis over a ferroptotic cell death.

LOPIT-DC data also supported the ferritin chains, FTH1 and FTL, as being differentially localized to the lysosome post-IR (Figure 5A-B). Confocal microscopy of ferritin and the lysosomal marker, lysosome-associated membrane glycoprotein 1 (LAMP1) was performed to validate the subcellular proteomic findings. Visual observations showed a dispersed staining of ferritin in the control samples with increased staining around the perinuclear area post-IR, which mimics the staining pattern seen by LAMP1 in both the control and IR-treated samples (Fig. 6A). Despite these visual observations, corresponding co-localization analysis was shown to not be significant (Fig. S10). Previous work using the A549 cell line has found significantly higher content of ferritin than a non-tumorigenic epithelial lung cell line counterpart and suggested intralysosomal ferritin provided a chelating function to maintain lysosomal membrane stability, preventing lysosomal leakage and subsequent autophagy, apoptosis, and necrosis (Persson et al., 2001). If autophagy is activated, ferritin is degraded by the lysosomes causing the cascade of cell death responses characteristic of ferroptosis (Gao et al., 2016; Hou et al., 2016). This process is known as ferritinophagy, which increases the bioavailability of iron in the cell, subsequently increasing sensitivity to ferroptosis (Jiang et al., 2021). Therefore, within the data presented here, it is difficult to determine whether the cells 12 hrs post-IR are undergoing active ferritinophagy, increasing ferroptosis sensitivity, or are chelating the endogenous iron to protect the cells from ferroptosis.

Additionally, in our LOPIT-DC data, TFRC was found to differentially localize from endoplasmic-reticulum–Golgi intermediate compartment (ERGIC) to the PM upon IR exposure (Fig. 5A-B) (Feng et al., 2020). Validation via imaging confirmed this with increased staining and co-localization of Membrite® and TFRC (p-value < 0.05) (Fig. 6B, D). Interestingly, control-TFRC appeared to have asymmetric perinuclear staining with this staining becoming more dispersed and evenly distributed around the perinucleus post-IR, similar to LAMP1 staining (Fig. 6A-B). This suggests ERGIC or Golgi apparatus localization within the control samples versus more active trafficking with potential lysosomal localization of TFRC post-IR. TFRC is a surface receptor, which internalizes transferrin-Fe^3+^ complexes via endocytosis and a significant regulator of intracellular iron. Increased intracellular iron is a key mediator of ferroptosis and increased expression of TFCR via MEK signaling in cancer cells have been attributed to higher sensitivity of some cancer cells to ferroptosis (Senyilmaz et al., 2015). Furthermore, PM-localized TFRC has recently been suggested as a reliable biomarker for ferroptosis, indicating that this observed increase in PM-targeting of TFRC could be sensitizing cells to ferroptosis (Feng et al., 2020). However, in our imaging data, the additional dispersed staining of TFRC visually appeared similar to LAMP1 staining post-IR and could also suggest an opposing anti-ferroptotic mechanism based on the following previous literature. In stress conditions, TFRC has been observed to shift its localization further down the endocytic pathway to the lysosomes. It is thought that TFRC is involved in recruitment of galectin 3 (LGALS3) to damaged lysosomes where it participates in ESCRT-III-mediated membrane repair and/or autophagy of damaged lysosomes. Lysosomal TFRC recruitment of LGALS3 is thought to favor ESCRT-III-mediated repair over autophagy (Jia et al., 2020a). This suggests a cell survival mechanism, rather than a pro-ferroptotic mechanism. To assess this, a further co-staining experiment was performed using LAMP1 as a lysosomal marker, but co-localization analysis of LAMP1 and TFRC post-IR showed no significance (Fig. S11). Further investigation with more optimized conditions is required to confirm this hypothesis.

Another observation with an interconnecting function with ferroptosis was the differential localization of hereditary hemochromatosis protein (HFE), which was found to localize to the PM after IR exposure (Fig. S8B). HFE is known to dimerize with TFRC at the PM to regulate iron intake by reducing TFRC affinity to transferrin (Feder et al., 1998; West et al., 2000). Generally, HFE has been studied in the context of hemochromatosis, where its most common mutation HFE (C282Y) prevents cell surface expression, destabilizing iron homeostasis and leading to iron overload in tissues, particularly the liver (Schmidt et al., 2008; West et al., 2000; Xiao et al., 2023). This finding suggests the cells are attempting to stabilize cellular iron levels and, therefore, an anti-ferroptotic response.

## Discussion

Within this study a combination of untargeted and targeted techniques were exploited to gain an increased insight of the expression and subcellular dynamics of proteins upon IR exposure, identifying 243 and 544 differentially-expressed and -localized proteins, respectively. Of these only 21 of the proteins measured were deemed to both change in abundance and subcellular localization, highlighting the complementary nature of these techniques and epitomizing the utility of using untargeted subcellular proteomics methods. Whilst proteins that changed in abundance or localization differed, the enriched pathways were highly interlinked, uncovering mechanisms such as iron metabolism and ECM organization.

Further targeted investigation into specific proteins emphasized subcellular-specific changes to a specific subset and behavior of proteins involved in ferroptosis-related signaling - ferritin (FTH and FTL1), TFRC and AIFM2. Lately, ferroptosis has come under the research spotlight, due to increased interest as a potential drug target for diseases such as cancer (Chen et al., 2023; Qi et al., 2022). This iron-dependent cell death response is characterized by induction of lipid peroxidation and iron overload. Our data suggests a prominent response to IR-induced lipid peroxidation and cellular iron homeostasis. Using these observations alongside prior literature, several hypotheses as to the function of these potential translocation events in relation to ferroptosis have been made (Fig. 7).

**Figure 7.**
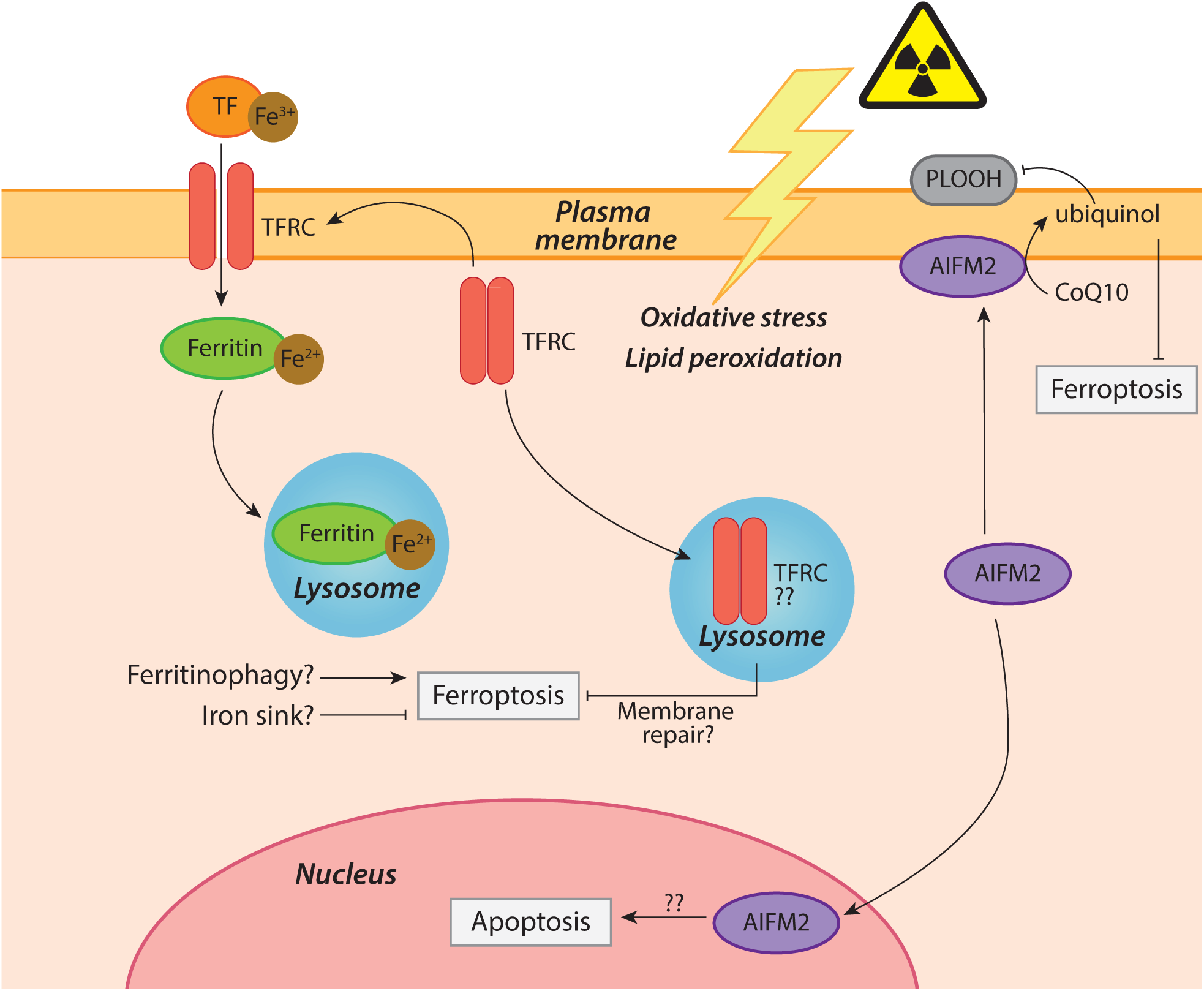
Schematic of key findings and potential implications within the ferroptotic pathways.

While PM-targeted TFRC suggested a pro-ferroptotic response to IR (Feng et al., 2020), the concurrent localization of HFE to the PM indicated dimerization of these two proteins and subsequent regulation and prevention of iron overload in the cells (Feder et al., 1998; West et al., 2000). Additionally, visual observations of the TFRC staining pattern in confocal microscopy appeared to mimic the distribution of LAMP1 staining. Lysosomal localization of TFRC has previously been suggested to participate in ESCRT-III-mediated membrane repair and therefore another protective mechanism (Jia et al., 2020a, 2020b). LOPIT-DC data showed both ferritin chains localized to the lysosomes post-IR. It has previously been shown that A549 cells have unusually high levels of ferritin compared with a non-tumorous counterpart and it has been hypothesized that the lysosomes are used as an iron sink to protect cells (Persson et al., 2001). This enhanced lysosomal integrity and lysosomal iron chelation capabilities could explain the challenge in treating lung tumors (Persson et al., 2005, 2005). However, lysosomal ferritin can also induce ferritinophagy, a ferroptoptosis-specific autophagy (Sun et al., 2022). Finally, AIFM2, also becoming more commonly known as ferroptosis suppressor protein 1 (FSP1), was found to localize to the PM upon IR. This is a characteristic translocation indicating suppression of ferroptosis via the antioxidant, glutathione-independent AIFM2/CoQ_10_ axis (Bersuker et al., 2019; Doll et al., 2019). Interestingly, proteins from the orthogonal glutathione-dependent cyst(e)ine/ GSH/GPX4 axis remained unchanged in both abundance and subcellular localization, indicating a potential preference or reliance of these cells to the AIFM2/CoQ_10_ axis for ferroptosis suppression.

A549 cells harbor KRAS mutation with RAS oncogenes have been implicated in radioresistance and found to be resistant to ferroptosis inducers, such as GPX4 inhibitors (Schott et al., 2015). Recent research has also shown elevated levels of AIFM2 in KRAS mutant tumors versus non-tumorous tissue, with elevated levels linked to tumour initiation and ferroptosis resistance (Müller et al., 2023). A549 cells are also kelch-like ECH associated protein 1 (KEAP1) mutants, which has recently been demonstrated to play a role in ferroptosis resistance via lack of inhibition of the AIFM2 transcription factor, nuclear factor erythroid 2-related factor 2 (NRF2). Inhibition of AIFM2 in combination with IR markedly increased radiosensitivity within these cells compared with IR or AIFM2i alone (Koppula et al., 2022).

These findings demonstrate the benefits of generating subcellular cell-wide maps for gaining insights into potential functions, particularly those with multifactorial purposes, which would otherwise be missed when looking at expression. The recent wave of new findings in the ferroptosis community, along with the findings in this study, solidifies the need for further research into the druggability of this cell death pathway in combination with current therapeutic options (Qi et al., 2022).

## Data availability

There is a dedicated R Shiny app for interactive viewing of the subcellular proteomics data online at http://proteome.shinyapps.io/a549lopit2024. All proteomics data have been deposited to the ProteomeXchange Consortium via the PRIDE partner repository (Perez-Riverol et al., 2022) with the dataset identifier PXD055123. Information related to the raw data files can be found in the Supplementary Data.

Reviewers can access the PRIDE entry via the following log-in details:

- Username:
- Password: 1yzsvjneMFai

## Acknowledgements

AstraZeneca for providing cell line, reagents and access to x-ray equipment. Mike Deery at the Cambridge Centre for Proteomics for mass spectrometry support. Joana Cerveira at Department of Pathology at the University of Cambridge for technical support and access to flow cytometer. Department of Pathology for use of their x-ray equipment. Nicola Lawrence and Richard Butler at the Gurdon Institute Imaging Facility for microscopy and image analysis support. Hilary J. Lewis and Ana J. Narvaez for project discussions. Soni Deshwal for comments on the manuscript. J.A.C was funded through a BBSRC iCASE award with Astra Zeneca (BB/R505304/1), L.M.B. by EU Horizon 2020 program INFRAIA project EPIC-XS (Project 823839), O.M.C by a Wellcome Trust Mathematical Genomics and Medicine studentship, K. S. L. by Wellcome Trust (110071/Z/15/Z) and European Union Horizon 2020 program INFRAIA project EPIC-XS (Project 823839).

## CRediT authorship contribution statement

**Josie A. Christopher:** Writing – original draft, Writing – review & editing, Investigation, Methodology, Visualization, Data curation, Formal analysis. **Lisa M. Breckels:** Writing – review & editing, Visualization, Software. **Oliver M. Crook:** Writing – review & editing, Visualization, Software. **Mercedes Vazquez--Chantada:** Methodology, Resources. **Derek Barratt:** Resources, Conceptualization, Funding acquisition. **Kathryn L. Lilley:** Methodology, Resources, Conceptualization, Funding acquisition, Project administration, Supervision, Writing – review & editing.

