## Supplementary figures for "Global proteomics indicates subcellular-specific anti-ferroptotic responses to ionizing radiation"

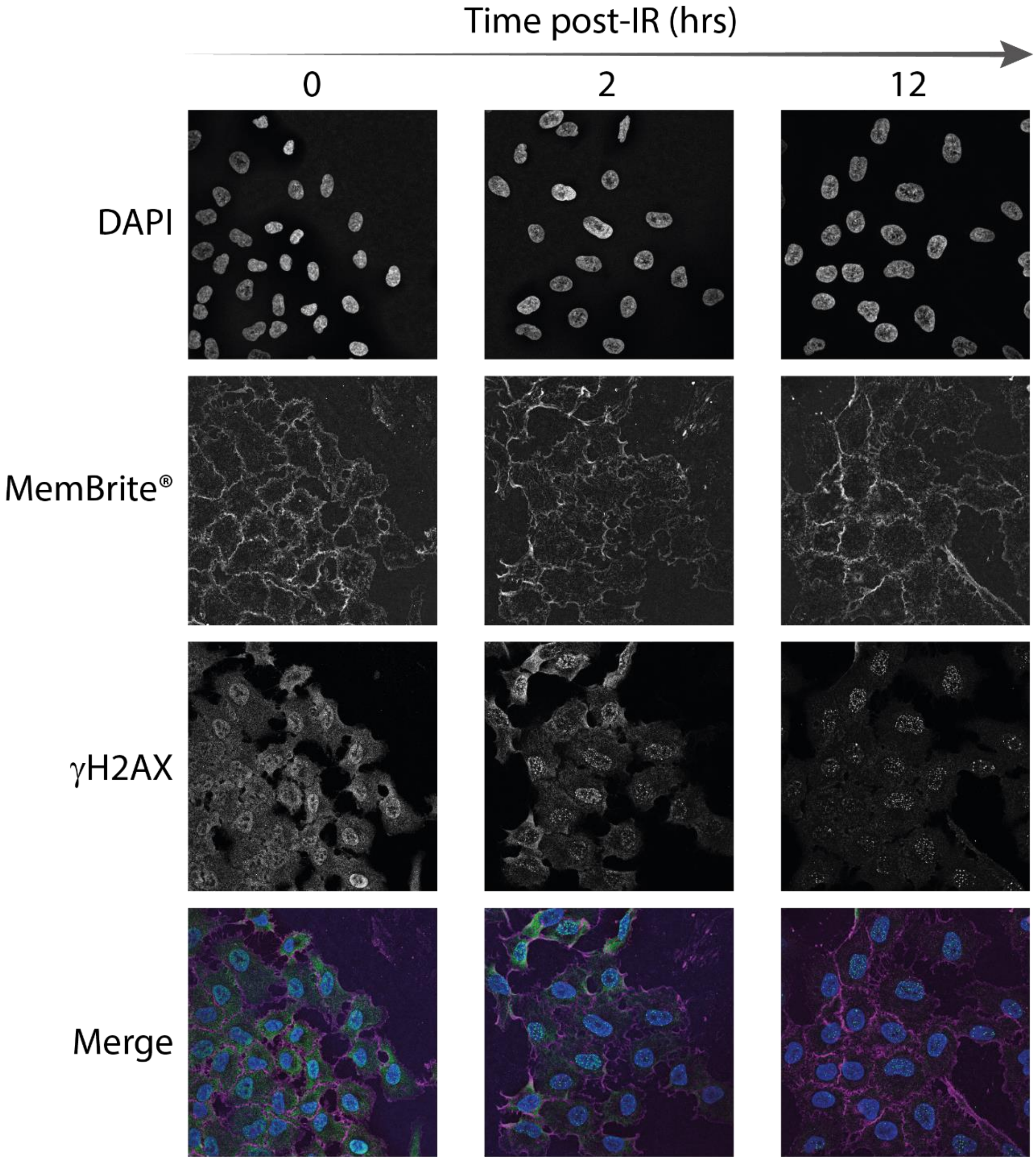


**Figure S1.** Examples of microscopy images used for measuring number of γH2AX nuclear and cytoplasmic foci at 0, 2 and 12 hrs post-IR. DAPI was used as a nuclear stain and MemBrite® as a PM stain to distinguish cell boundaries. Colour image shows all three channels merged into one image (DAPI: blue, MemBrite®: magenta, γH2AX: green).

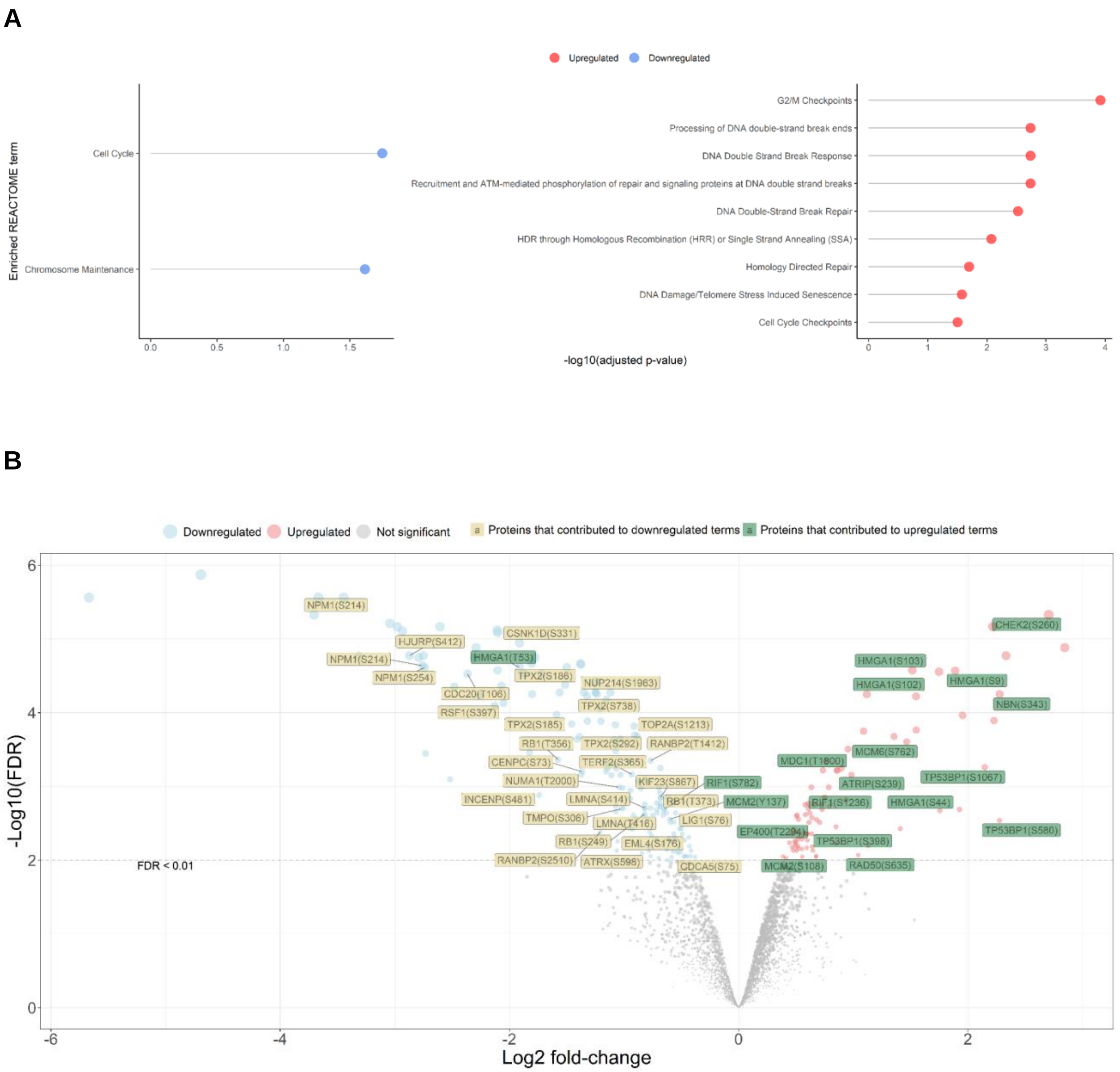


**Figure S2.** Corresponding phosphoproteomics data of same samples that underwent total proteomics analysis. (A) Enriched REACTOME terms for the proteins the up- and down-regulated phosphopeptides derive from at 12 hrs post-IR, and (B) these phosphosites annotated on volcano plot from differential expression analysis of phosphoproteomics data.

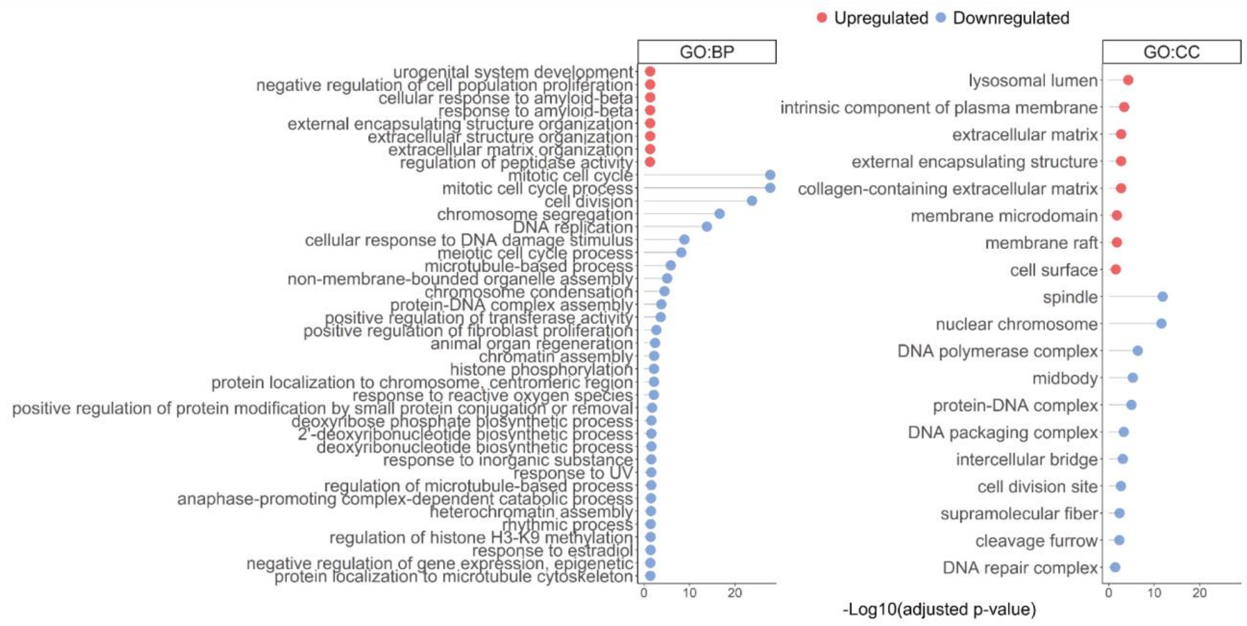


**Figure S3.** Enriched biological process (BP) and cellular component (CC) gene ontology terms for down- and up-regulated proteins (FDR < 0.01) 12 hrs post-IR in the total proteomics data.

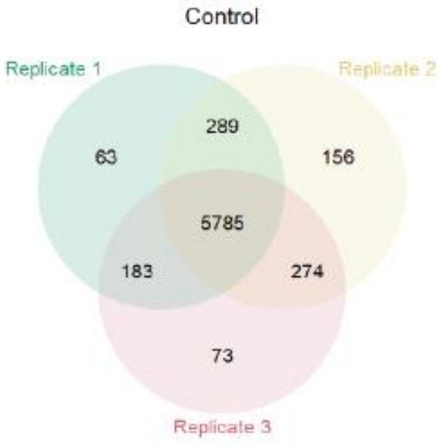


**Figure S4.** Venn diagrams demonstrating overlap of protein identifications between LOPIT-DC replicates in control and 12 hrs post-IR group.

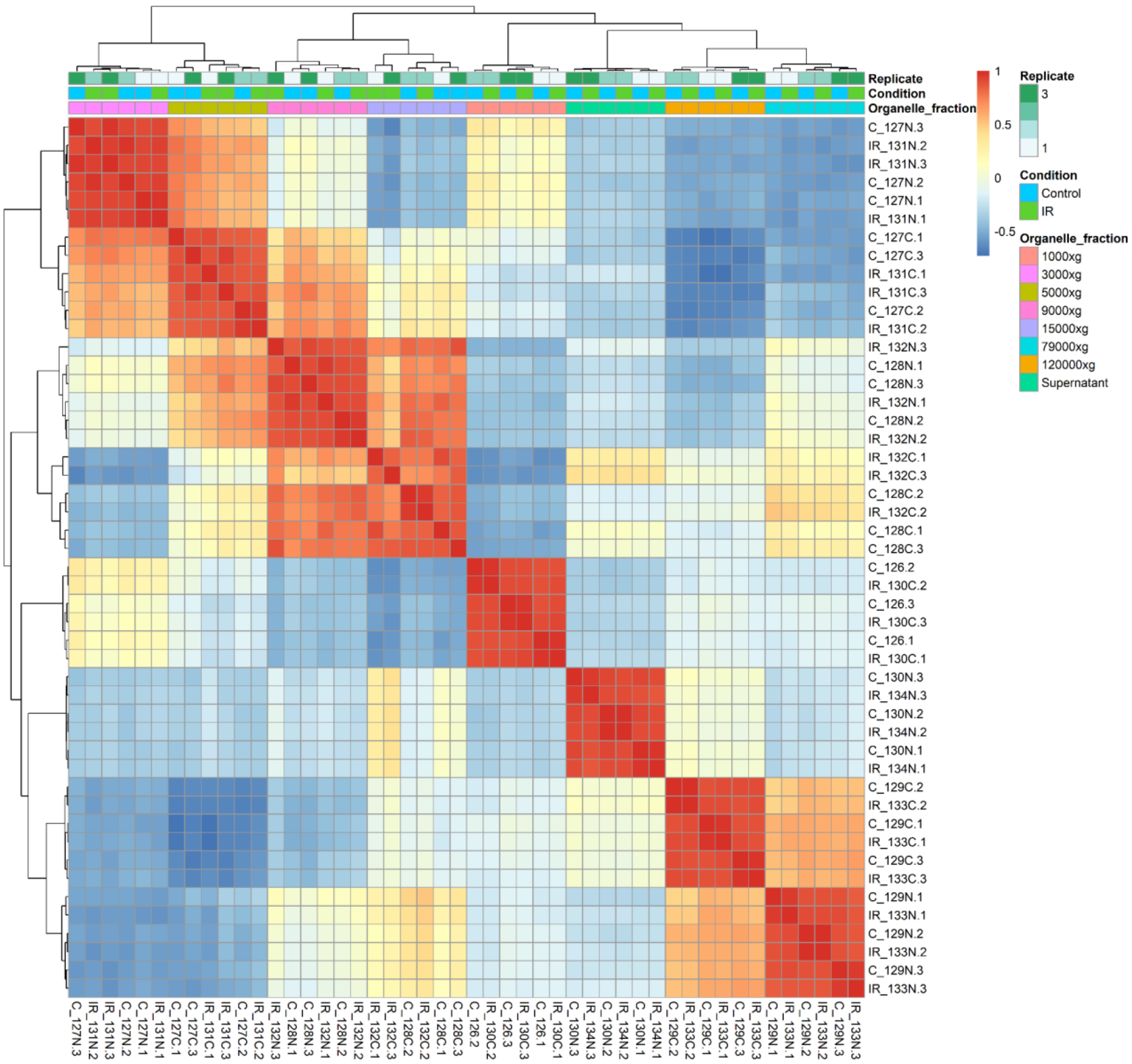


**Figure S5.** Pair-wise Pearson's correlation matrix for each organellar lysate fraction for each LOPIT-DC experiment.

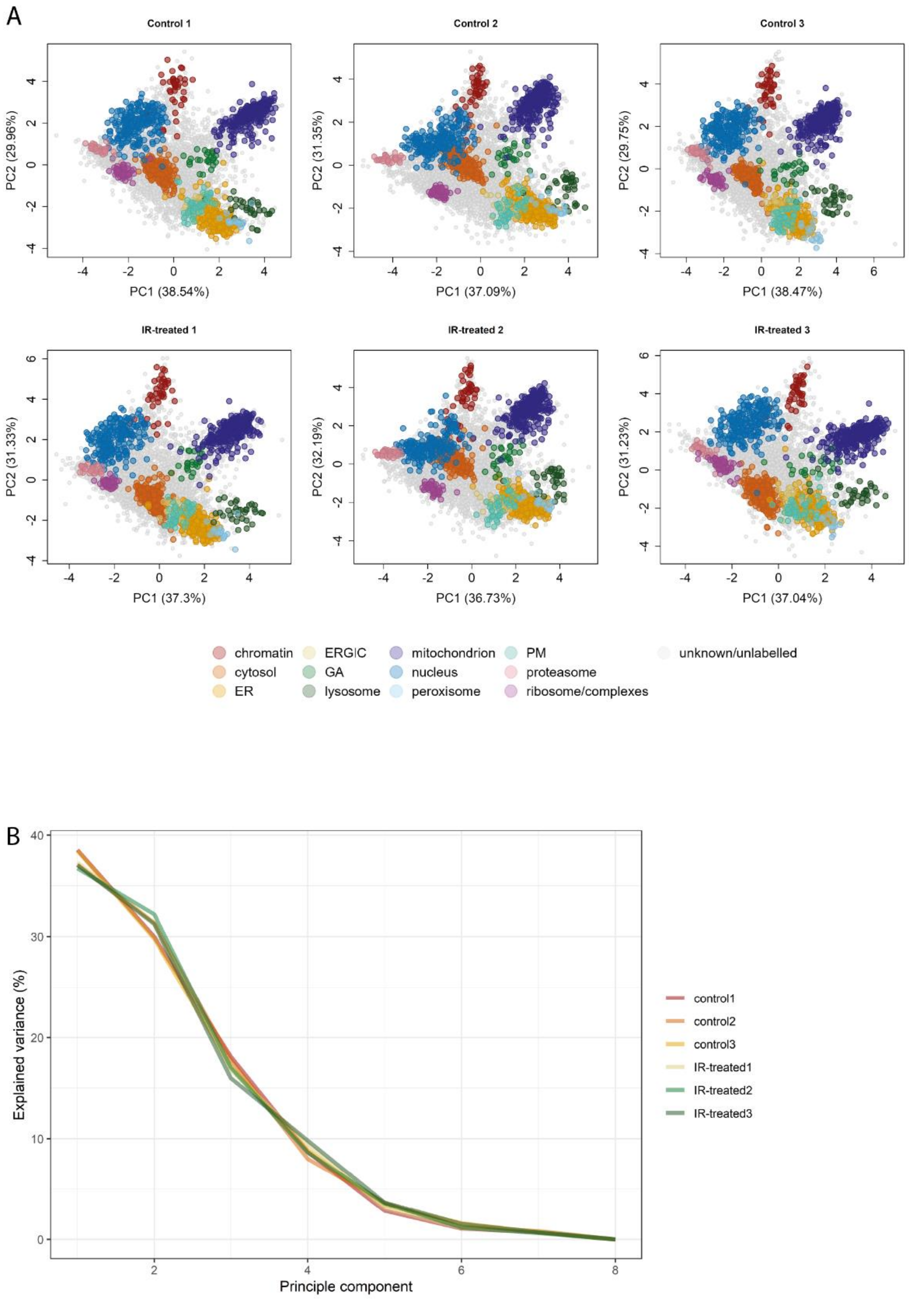


**Figure S6.** (A) Principle component analysis (PCA) plots for individual LOPIT-DC replicates. Each colour denotes the protein markers of 12 distinct subcellular compartments. (B) Corresponding skree plot showing percentage of explained variance across each principle component with each colour representing an individual LOPIT-DC replicate. (ER, endoplasmic reticulum; ERGIC, endoplasmic-reticulum–Golgi intermediate compartment; GA, Golgi apparatus; PM, plasma membrane).

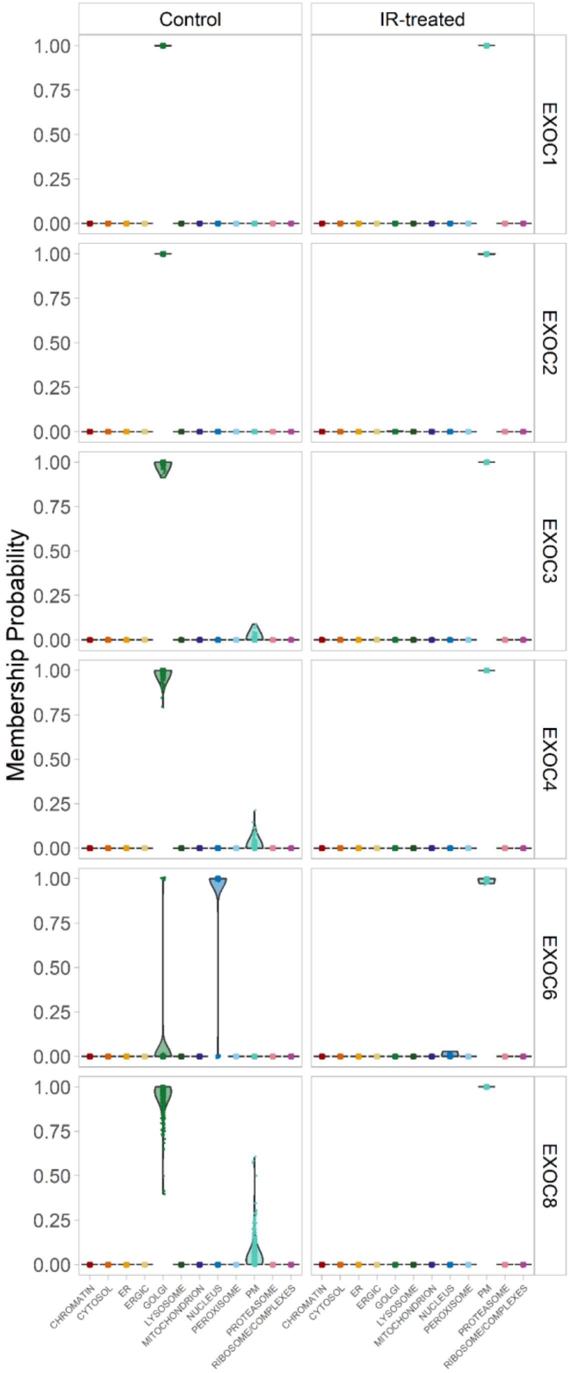


**Figure S7.** Posterior predictive distributions (membership probability) across the 12 subceullar compartments for proteins identified as differentially localised and part of the exocyst complex in control vs IR-treated LOPIT-DC samples. (ER, endoplasmic reticulum; ERGIC, endoplasmic-reticulum–Golgi intermediate compartment; GA, Golgi apparatus; PM, plasma membrane).

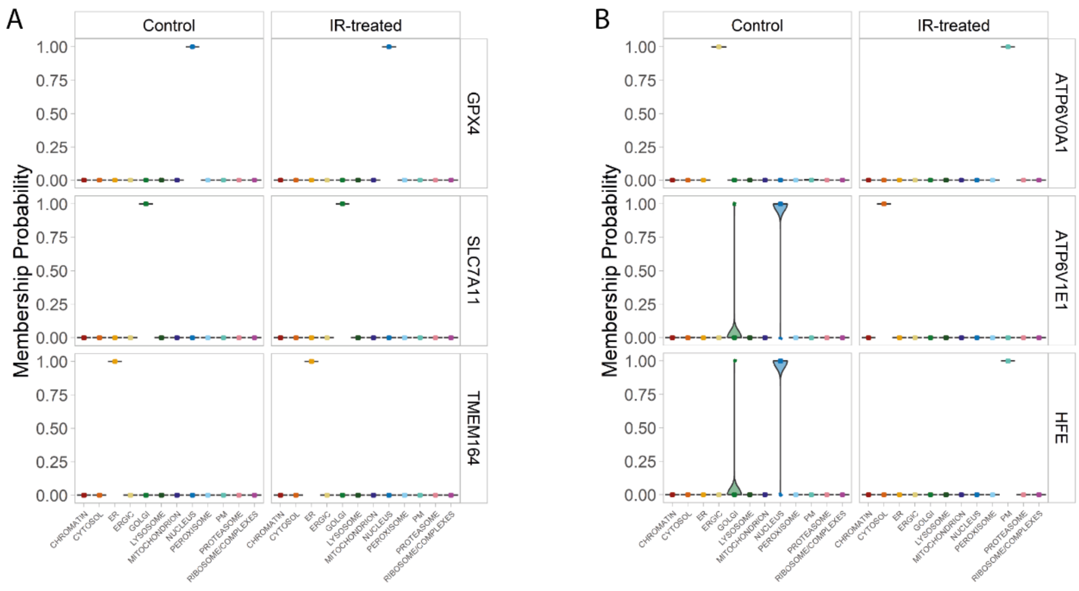


**Figure S8.** Posterior predictive distributions (membership probability) across the 12 subceullar compartments for proteins identified as (A) not differentially localised and part of ferroptosis pathway, and (B) differentially localised and part of iron uptake and transport in control vs IR-treated LOPIT-DC samples. (ER, endoplasmic reticulum; ERGIC, endoplasmic-reticulum–Golgi intermediate compartment; GA, Golgi apparatus; PM, plasma membrane).

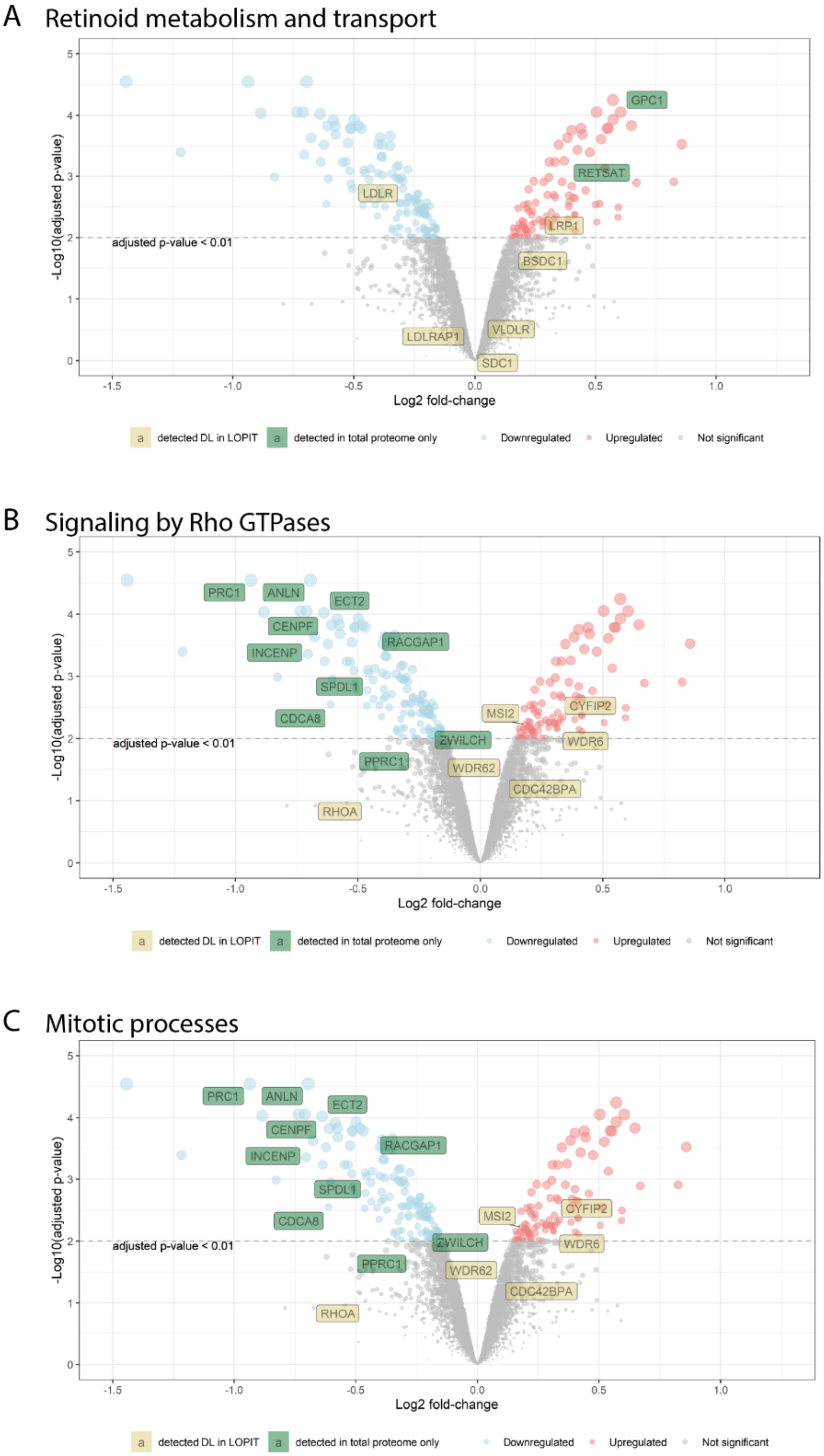

A Retinoid metabolism and transport
adjusted p-value < 0.01
-Log10(adjusted p-value)
Log2 fold-change
GPC1
RETSAT
LDLR
LRP1
BSDC1
VLDLR
LDLRAP1
SDC1
detected DL in LOPIT
detected in total proteome only
Downregulated
Upregulated
Not significant
B Signaling by Rho GTPases
adjusted p-value < 0.01
-Log10(adjusted p-value)
Log2 fold-change
PRC1
ANLN
ECT2
CENPF
RACGAP1
INCENP
SPDL1
CDCA8
MSI2
CYFIP2
ZWILCH
WDR6
PPRC1
WDR62
CDC42BPA
RHOA
detected DL in LOPIT
detected in total proteome only
Downregulated
Upregulated
Not significant
C Mitotic processes
adjusted p-value < 0.01
-Log10(adjusted p-value)
Log2 fold-change
PRC1
ANLN
ECT2
CENPF
RACGAP1
INCENP
SPDL1
CDCA8
MSI2
CYFIP2
ZWILCH
WDR6
PPRC1
WDR62
CDC42BPA
RHOA
detected DL in LOPIT
detected in total proteome only
Downregulated
Upregulated
Not significant

**Figure S9.** Volcano plot of differential expression analysis for the total proteomics data in control verus 12 hrs post-IR in A549 cells annotated with proteins involved in (A) retinoid metabolism and transport, (B) signalling of Rho GTPases, and (C) mitotic processes. Coloured points denote all proteins with FDR < 0.01 and coloured labels denote proteins detected as differentially localised (DL) in LOPIT-DC (yellow) or only detected in total proteomics (green).

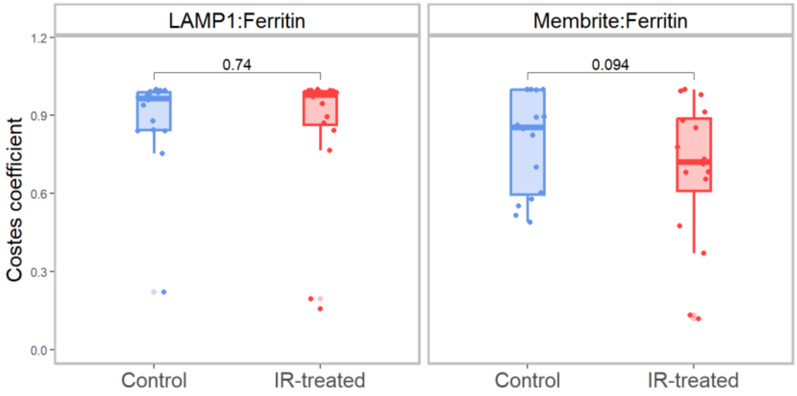


**Figure S10.** Quantitative microscopy analysis to assess co-localisation of ferritin and LAMP1 immostaining, and Membrite® staining.

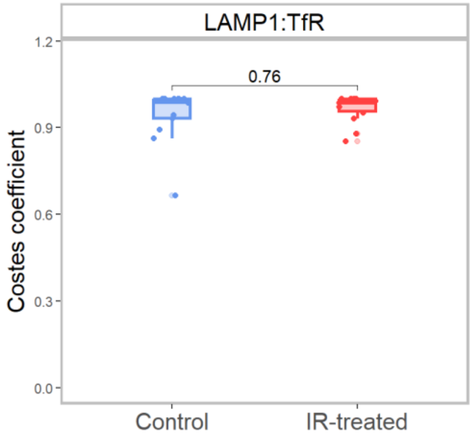


**Figure S11.** Quantitative microscopy analysis to assess co-localisation of LAMP1 and TFRC (TfR).
