## Supplementary methods for "Global proteomics indicates subcellular-specific anti-ferroptotic responses to ionizing radiation"

### Experimental Procedures

#### Cell culture

Lung epithelial carcinoma cell line, A549 (CCL-185™, ATCC®) were certified as mycoplasma-free. Cells were not kept in culture for longer than a month and were grown in Ham's F12 (Sigma Aldrich), with 10% fetal bovine serum (FBS, ThermoFisher, Lot: 42F3393K) and 1% L-glutamine. Cells were passaged at around 70% confluence by washing with PBS before dissociating from the flask with TrypLE™ Express (Gibco™, 12604013) and kept at 37°C and 5% CO2.

#### X-ray treatment

Total absorbed dose of 6 Gy using either a Faxitron® CellRad® or Pantak HF320kV x-ray system. The Pantak HF320kV x-ray system operated at 220 kV, 14 mA with a 0.5 Cu filter with a dose rate of 428 cGy/min. The Faxitron® CellRad® using a 50 Cu with a kV/mA dependent on the shelf used on the instrument. Control cells were sham irradiated (kept in the same conditions as the IR-treated cells).

#### Localisation of Proteins using Isobaric Tagging using Differential Centrifugation (LOPIT-DC)

Three T175 flasks at approximately 90% confluence were used for each LOPIT-DC sample (equating to ~7,000 µg protein). Three biological replicates were performed. Cells were harvested using TypeLE™ Express at 37°C for 7 mins. Cell suspensions were then transferred to Falcon tubes and washed twice using chilled PBS and spinning the cells at 200 xg for 5 mins at 4°C. Resulting cell pellet was resuspended in an isotonic lysis buffer (250 mM sucrose, 10 mM HEPES (pH 7.4), 2 mM EDTA (pH 8.0), 2 mM magnesium acetate tetrahydrate, supplemented with cOmplete™ protease and ROCHE PhosSTOP™ phosphatase inhibitors (Sigma-Aldrich, 11836170001, 4906845001). Twelve hours after (sham) irradiation, cell suspensions were then passed through a ball-bearing homogeniser (Isobiotec) using a 12 µM clearance 25 times to achieve ~90% lysis, assessed via Trypan blue staining and a haemocytometer.

Differential centrifugation was performed using a combination of a refrigerated benchtop centrifuge (Eppendorf 5415R) and a Beckman Optima MAX-XP ultracentrifuge (with a TLA-55, fixed angle rotor) with spin speeds and times in Table S1 (Fig. 1). Samples were always kept on ice or at 4°C to maintain organelle integrity. The first spin was used to clear unlysed cells from the sample and discarded. The proteins in the final supernatant were precipitated using at least four-fold volume of chilled acetone and stored at -20°C overnight, before pelleting at top speed for 20 mins at 10°C and left to air dry for 15 mins. Organellar-enriched pellets were resuspended in solubilisation buffer (8 M urea, 0.2% SDS, 50 mM HEPES (pH 8.5)) and sonicated for 15 mins at 30 s (on/off) intervals using a Diagenode Bioruptor® Plus with water cooling system. Fractions were then spun at 16,000 xg for 5 min to assess whether pellets had solubilised. If pellets were visible, further solubilisation buffer was added and sonication repeated. Organellar enriched lysates then underwent reduction, alkylation, digestion, TMT labelling, clean-up and LC-MS/MS as documented below.

#### Total & phospho-proteomics sample harvesting & lysis

Two T75 flasks at approximately 90% confluence were used per sample (equating to ~1,200-1,400 µg protein). Three biological replicates were performed. Cells were washed twice with PBS before dissociation with TrypLE™ Express for 5 mins at 37°C, then washed twice using chilled PBS and spinning at 200 xg for 5 mins at 4°C. Twelve hours after (sham) irradiation, cell pellets were then resuspended in lysis buffer (8 M urea, 0.2% SDS, 50 mM HEPES (pH 8.5)) supplemented with protease and phosphatase inhibitors) and sonicated using a Vibra-Cell probe sonicator (Sonics) at 50 joules for three cycled with 10 sec pauses between cycles to prevent overheating of the sample.

#### Protein digestion, TMT labelling & clean-up

Protein concentrations in LOPIT-DC pellets and total proteomics samples were measured using BCA protein assay (Pierce™, 23225) according to the manufacturer's microplate protocol. Absorbance was measured on a Multiskan SkyHigh Microplate Spectrophotometer (ThermoFisher) at a wavelength of 562 nm using the precision setting and a four-parameter quadratic standard curve. Any measurements with CVs of >5% were repeated. Samples with poor measurements due to viscosity from DNA were were incubated at room temperature for 20 mins with nuclease enzyme (Millipore Benzonase® Nuclease HC; 71206-3). Buffer blanks were used and if interfering substances were present in the buffer, the samples were diluted to below recommended thresholds. Samples and pellets were normalised to 50 µg in 100 µLlysis buffer each.

Total proteomics samples and LOPIT-DC pellets were reduced with 10 mM dithiothreitol (DTT) for 1 hr at 37°C, followed by alkylation with 25 mM iodoacetamide (IAA) for 2 hrs at room temperature in the dark. Suspensions were then precipitated using acetone (as above) and pellets resuspended in 100 mM HEPES (pH 8.5) before sonication for 15 min with 30 s on/off intervals. Two-step protein digestion was performed using sequencing-grade trypsin (Promega, V5111) at a 1:40 enzyme:protein ratio for

1 hr at room temperature on a shaker, followed by adding the same amount to give a total of 1:20 enzyme:protein ratio before overnight incubation at 37°C.

Tandem mass tag (TMT) labelling was performed as per the manufacturer's instructions. TMT 10-plex (ThermoFisher, 90406) were used for total proteomics sample labelling and TMTpro 16-plex (ThermoFisher, A44520 (pre-release batch)) were used for LOPIT-DC pellet labelling. As TMT labels are designed to label 100 µg protein, each TMT label reagent was split into two to label 50 µg protein. Samples were lypholised using a SpeedVac (Labconco).

Solid phase extraction performed using SepPak® C18 columns (Waters, WAT054955). Briefly, columns were conditioned with 100% ACN, followed by 0.05% acetic acid (pH 3), followed by 0.1% TFA. Samples were resuspended and loaded into columns using 0.1% TFA (pH 2) followed by three 0.1% TFA washes and two 0.05% acetic acid washes. Peptides were eluted with 70% ACN/0.05% acetic acid and lyophilized.

#### Phosphopeptide enrichment

Following solid phase extraction, dried peptides were resuspended in 80% ACN/5% TFA/1 M glycolic acid and phosphopeptides enriched using $TiO_2$ beads (GL Sciences cat. no. 5020-75010) at a concentration of 0.6 mg beads per 100 µg peptides. After 15 min on shaker, beads were pelleted and supernatant underwent another round of $TiO_2$ enrichment with half the concentration of beads and then again for a third round of $TiO_2$ enrichment. The three resulting $TiO_2$ bead pellets were pooled and washed with 80% ACN/1% TFA, followed by an additional wash with 10% ACN/0.1% TFA and lyophilized. Phosphopeptides were eluted with 25% ammonia solution (pH 11.3) by shaking the beads for 15 min, beads were pelleted, and supernatant passed through a C8 stage tip (in-house made). Finally, 30% ACN was used to elute any bound peptide in the C8 material and added to the phosphopeptide enriched pool. Peptides were immediately lyophilized. All incubation steps were performed at room temperature.

#### Pre-fractionation

Total proteomics and LOPIT-DC samples were pre-fractionated offline using reverse phase UPLC with an Acquity UPLC System with diode array detector (Waters). Peptides were loaded onto an Acquity UPLC BEH 283 C18 column (2.1-mm ID × 150- mm; 1.7-µm particle size) (Water, 186002353) using an aqueous mobile phase 20 mM ammonium formate (pH 10) buffer. Peptides were eluted across a 50 min linear gradient from 5 to 75% ACN/20 mM ammonium formate (pH 10) at a flow rate of 0.24

mL/min. Peptide fractions were eluted into separate tubes every minute of the gradient. This resulted in 35 fractions that were concatenated into 15 fractions. Samples were lyophilised and stored at -20$^{o}$C until MS analysis.

Phosphoproteomics samples were pre-fractionated offline using the Pierce$^{TM}$ high pH reverse-phase peptide fractionation kit (ThermoFisher, 84868) according to manufacturer's protocol.

## LC-MS/MS

Dried samples were resuspended in 0.1% formic acid at an estimated concentration of 0.5 μg/μL and injection volumes dependent on achieving an estimated 1-1.5 μg of peptide. Analysis was performed on a Lumos Orbitrap mass spectrometer (Thermo Fisher Scientific Inc, Waltham, MA, USA) coupled to a Dionex Ultimate 3000 RSLC nanoUPLC (Thermo Fisher Scientific Inc, Waltham, MA, USA) system for online pre-fractionation. Samples were loaded onto the pre-column (Thermo Scientific PepMap 100 C18, 5mm particle size, 100Å pore size, 300 mm i.d. x 5mm length) via the Ultimate 3000 auto-sampler for 3 min at a flow rate of 15 µL/min. Reverse-phase chromatography was performed with a nanoEasy-spray column (Thermo Scientific PepMap C18, 2mm particle size, 100 Å pore size, 75 mm i.d. x 50 cm length) at a flow rate of 300 nL/min. The mobile phase during loading was 0.1% formic acid with a linear gradient of 2-40% of 80% ACN/0.1% formic acid over 93 min (with a total run time of 120 min inclusive of high organic wash steps and column equilibration). Eluted peptides were injected into the MS analyser via electrospray ionization (ESI) using an Easy-Spray source (Thermo Fisher Scientific). Peptide m/z values were measured in the Orbitrap analyser, with a resolution of 120,000 with a 380-1,500 Da m/z range. Data-dependent MS/MS (MS2) scans using Top Speed setting were used to isolate and fragment precursor peptide ions from MS1 using CID (NCE: 35%) and were analyzed in the linear ion trap (total and subcellular proteomics) or in the Orbitrap analyser with resolution of 30,000 (phosphoproteomics). Peptides with single or unassigned charge states were excluded from selection from MS1 and a dynamic exclusion window of 70 seconds was employed. Synchronous Precursor Selection (SPS)-MS3 was utilized to select the top 10 most abundant fragment ions for subsequent further fragmentation with HCD (NCE: 65%) within the high-energy collision cell. TMT reporter ion m/z values and relative abundances were measured in the Orbitrap analyser within the mass range of 100-500 Da at a resolution of 50,000 (total and subcellular proteomics) or 60,000 (phosphoproteomics). Cycles of 10 MS3 events occur before the instrument starts a new MS1 scan.

#### Flow cytometry

Two wells of a 6-well plate were prepared per sample for the cell death and cell cycle flow cytometry assays. Cells were dissociated with TrypLE™ Express for 5 mins at 37°C and were washed twice in chilled PBS and spinning at 200 xg for 5 mins at 4°C. Manufacturer protocols for propidium iodide (Abcam, ab139418) and CellEvent™ Caspase-3/7 (Invitrogen™, C10427) staining were followed for the cell cycle and cell death assays, respectively. Cells were treated with 1 μg/ml nocodazole for 18 hrs (as previously published (Queiroz et al., 2019)) or 100 μg/mL sodium arsenate for 2 hrs as positive controls for cell arrest and cell death, respectively. Sodium arsenate was also added to compensation samples. Cells were analysed on an Attune™ NxT Acoustic Focusing Cytometer using the yellow-green laser (561 nm) and the YL2 bypass filter (620/15); and the blue (488 nm) laser with the BL1 (530/30) and BL2 (695/40) dichroic filters, for the cell cycle and the cell death assay, respectively. Both experiments used a flow rate of 100 μL/min and set to collect 100,000 cell events.

#### Western blotting

Samples were run on Bio-Rad® mini-PROTEAN® TGXTM pre-cast gels (4-20% or 7.5% polyacrylamide) in BioRad® Tris-glycine-saline (TGS) solution on a Bio-RADTM Criterion™ Vertical Electrophoresis Cell using a Bio-Rad® PowerPacTM 1000 power supply. Laemmli buffer was added to samples before loading with 30 μg for whole cell lysates and run alongside a protein standard (BioLab® Dual Colour 11-245 kDa). Gels were transferred to Bio-Rad® TransBlot® 0.2 μm PVDF membrane on "Turbo" setting using a Bio-Rad® TurboBlot® system and blocked for 1 hr at room temperature in 5% milk in Tris buffer solution with Tween (TBST). Membranes were probed overnight at 4°C with 1:1000 dilution of anti-yH2AX (Abcam, ab22551) in 5% milk in TBST and then washed three times in TBST. Secondary HRP-linked antibodies (Amersham, NA934, NA931) at 1:10,000 dilution in 5% milk in TBST were incubated for 2 hrs at room temperature. Membranes were washed three times with TBST before applying enhanced chemiluminescence (ECL) (Bio-Rad, 1705061) reagent and developed onto x-ray film in a dark room.

#### Immunofluorescence microscopy

Cells were seeded into a glass-bottomed 8-high-well μ-Slide (Ibidi, 80806) at approximately 2-11x$10^4$ cells/mL. At ~60-70% confluence, slides were exposed to 6 Gy x-ray and incubated for 12 hrs. Cells were stained with MemBrite® Fix 640/60 cell surface stain (Biotium, 30097) prior to fixation. Staining was performed as according to manufacturer's guidelines. Cells were then washed with Hank's Balanced Salt Solution (HBSS, ThermoFisher, 24020091) three times prior to fixing with 4%

paraformaldehyde (PFA) in PBS and incubated for 15 min at room temperature. PFA was then removed and cells washed three times with PBS. Cells were then permeabilised using 0.1% Triton X-100 in PBS and incubated for 5 min at room temperature and washed three times with PBS. Primary antibodies were then added at the manufacturer or optimised concentration and incubated overnight at 4$^{o}$C. The following day, cells were washed four times with PBS and incubated with corresponding IgG (H+L) conjugated with Alexa Fluor 568 (Invitrogen, A11004) or Alexa Fluor 488 (Invitrogen, A-11008) with 1:2,000 dilution. Then washed four times with PBS and mounted using ProLong™ Gold Antifade Mountant with DAPI (Invitrogen, P36931). Single-stained and secondary only controls were also prepared to assess non-specific binding or autofluorescence. See Table S2 for antibody details.

A Zeiss LSM 880 confocal microscope with a 40x/1.30 Plan Apo Oil objective lens and a Piezo stage was used for image acquisition. Fluorophores were excited using 405 nm (blue diode), 488 nm (Argon), 561 nm (He 543) and 633 nm (He 633) lasers with 0.05%, 1.5% and 3.0% power, respectively. Signal was detected using two photomultiplier (PMT) and a gallium arsenide phosphide (GaAsP) array detector in combination with filters set to 400-440 nm, 507-552 nm, 585-620 nm, 650-690 nm, to detect DAPI, AlexaFluor488, AlexaFluor568, and MemBrite® Fix 640/60, respectively. Master gain was adjusted accordingly to prevent signal saturation in the detectors (target-specific), digital gain set to 1 and digital offset set to 0. The pinhole was adjusted according to the largest target wavelength (660 nm), giving a pinhole size of 44 μm. All collected images were 2048 x 2048 pixels with a pixel dwell of 4.10 μs.

#### MS data processing

Raw MS files were processed using Proteome Discoverer 2.4 (ThermoFisher) and a Mascot server (Matrix Science) v2.6 (Brosch et al., 2009). LOPIT-DC samples were analysed using the same processing and multi-consensus workflow against a SwissProt *Homo sapiens* database (no isoforms, 20,245 sequences, downloaded 09/04/2018) and the common repository of adventitious proteins (cRAP, v1.0, 123 sequences). An MS1 and MS2 ion tolerance of ±15 ppm and ±0.6 Da for protein identifications was used, respectively. Allowance of 2 missed tryptic cleavages and a minimum of 2 unique peptides per protein identification were set. An interference/co-isolation threshold of 50 with no signal:noise filtering was applied. Fixed modifications were set to carbamidomethyl(C) and TMT 10-plex or TMTpro(K, peptide N termini), and variable modifications were set to carbamyl(N-term), carbamyl(K), carbamyl(R) and deamidation(N,Q). Percolator was used to assess the false discovery rate (FDR) and only peptides with FDR $< 0.01$ were retained. Additional data reduction filters were: peptide rank = 1 and ion score >20. Quantification of the reporter ions calculated using centroided ions within a ±2 millimass unit (mmu) around the expected m/z of each TMT reporter ion and batch-specific TMT reporter ion isotope distributions were accounted for. TMT labelling efficiency was checked by using the same parameters, except setting TMT 10-plex or TMTpro as a variable modification.

#### Data analysis - flow cytometry

All flow cytometry analysis was performed using the FCS ExpressTM software (De Novo Software™, version 7.12). Gates were set to exclude debris and doublets from the analysis. Cell cycle analysis was performed using the MultiCycle algorithm feature in the FCS Express software, using the AutoFit function (1 cycle, Model 1, Dean & Jett mathematical model (Dean and Jett, 1974)). Compensation matrixes were set using the compensation control samples prepared alongside the experimental samples. Statistical analysis and visualisation for both the cell cycle and cell death assay results were performed in R using a pair-wise t-test and Bonferroni adjustment.

#### Data analysis - proteomics

Peptide spectrum match (PSM)-level quantitation output from Proteome Discoverer was uploaded into R. Experimental contaminants (identified with cRAP database), PSMs with no quantitation information, non-unique peptides and PSMs with no accession number were filtered from the datasets. Proteins with a signal:noise ratio of <10 were also filtered out based on previously found artifact with TMT quantitation using Tune v2 MS software (Hughes et al., 2017; Smith et al., 2022). All proteins with missing values were filtered out. PSMs were aggregated to proteins using the "robust" method within the pRoloc R package (v 1.34.0). The total proteomics dataset was normalised using (variance stabilising normalisation) (VSN) and LOPIT-DC datasets sum normalised. Replicates from LOPIT-DC datasets were combined using the combine function in pRoloc to generate a single subcellular map per experimental condition. The Limma R package was used to perform empirical Bayes moderated t-test (Smyth, 2004). P-values were adjusted for multiple testing (Benjamini and Hochberg, 1995). GO enrichment analysis was performed by using the Gene Ontology PANTHER search engine or using the R package ClusterProfiler (Gene Ontology Consortium, 2021; Wu et al., 2021).

#### Data analysis - BANDLE

A semi-supervised machine learning approach, Bayesian ANalysis of Differential Localisation Experiments (BANDLE), was used to classify proteins to subcellular localisations, identify differential localisation, and estimate uncertainty of those predictions (Crook et al., 2021). Subcellular protein markers for this algorithm were established using existing curated markers (Geladaki et al., 2019; Mulvey et al., 2017; Orre et al., 2019), the bioinformatics tool COMPARTMENTS (Binder et al., 2014), and the unsupervised, clustering algorithm, HDBscan (Melvin et al., 2016). The proteins selected as marker candidates using these methods were then filtered based on how reproducible their data was across the LOPIT-DC datasets by calculating distance correlation (those with distance correlation of >0.8 were selected). Final filtering of markers was performed visually by assessing the protein correlation profiles and PCA plots, as well as literature evidence. BANDLE analysis was performed as documented in Bioconductor vignettes (https://bioconductor.org/packages/release/bioc/html/bandle.html) with hyperparamters set to 10, 60 and 100 for length-scale, amplitude, variance, respectively. The estimated FDR (eFDR) was calculated as described in the vignette and a differential localisation score corresponding with a eFDR < 0.05 cut-off (0.57) was used to distinguish differentially localised proteins.

#### Data analysis - Microscopy

Segmentation and quantification of nuclear-bound γH2AX foci was performed using the open-source CellProfiler software (v4.2.1. Broad Institute) (Stirling et al., 2021). The same parameters were used from the CellProfiler "speckle analysis" tutorial. Briefly, images were uploaded as CZI files, grouped into experimental conditions, separated into separate channels and converted to grayscale images. Nuclei were then segmented based on a typical diameter of 100-300 pixels using a two-class, global Otsu threshold. Declumping method was set based on shape. γH2AX foci were enhanced prior to segmentation using the "enhance speckles" function using a feature size of 8 and the "slow" setting. Foci were then masked to only include foci that land within nuclei before proceeding to foci segmentation. Masked foci were segmented using a typical diameter of 4-20 pixels with global robust background thresholding (lower/upper outlier fraction of 0.05) accounting for mean and variance (standard deviation of 2.0). A threshold smoothing scale of 1.3488 and correction factor of 1 was used (lower/upper bounds of threshold 0 and 1). Declumping of foci was based on intensity. Data was collected on object intensity and relationships between daughter and parental objects (i.e. number of foci within individual nuclei). A similar process was applied to quantify cytosol-bound γH2AX foci. Co-localisation analysis was also performed using CellProfiler. Briefly, intensities for each channel were scaled and log2 transformed before applying a median filter and measuring Costes across entire image using the "Faster" thresholding method.
