## Supplementary tables for "Global proteomics indicates subcellular-specific anti-ferroptotic responses to ionizing radiation"

**Table S1.** Spin speeds and duration of each centrifugation within the differential centrifugation steps of the LOPIT-DC workflow.

| Fraction | Speed (xg) | Time (min) |
|---|---|---|
| Unlysed cells | 200 | 5 |
| Pellet 1 | 1,000 | 10 |
| Pellet 2 | 3,000 | 10 |
| Pellet 3 | 5,000 | 10 |
| Pellet 4 | 9,000 | 15 |
| Pellet 5 | 15,000 | 15 |
| Pellet 6 | 79,000 | 43 |
| Pellet 7 | 120,000 | 45 |
| Supernatant | - | - |

**Table S2.** Primary antibodies used in immunofluorescence microscopy experiment.

| Target | Gene name | Product number | Manufacturer | Species | Used dilution/ concentration |
|---|---|---|---|---|---|
| Ferroptosis suppressor protein 1 | AIFM2 | ab219986 | Abcam | rabbit | 0.25-2 μg/mL |
| Transferring receptor 1 | TFRC | ab84036 | Abcam | rabbit | 5 μg/mL |
| Ferritin | FTH1/FTL | ab75973 | Abcam | rabbit | 1:100 |
| Gamma-histone H2AX | γH2AX | ab22551 | Abcam | mouse | 2-4 μg/mL |
| Lysosome-associated membrane glycoprotein 1 | LAMP1 | ab25630 | Abcam | mouse | 5-10 μg/mL |
